# Assembly of a functional neuronal circuit in embryos of an ancestral metazoan is influenced by environmental signals including the microbiome

**DOI:** 10.1101/2024.12.10.627692

**Authors:** Christopher Noack, Sebastian Jenderny, Christoph Giez, Ornina Merza, Lisa-Marie Hofacker, Jörg Wittlieb, Urska Repnik, Marc Bramkamp, Karlheinz Ochs, Thomas C. G. Bosch

## Abstract

Understanding how neural populations evolve to give rise to behavior is a major goal in neuroscience. However, the complexity of the nervous system in most invertebrates and vertebrates complicates the deciphering of underlying fundamental processes. Here, we explore the self-assembly of neural circuits in *Hydra*, an organism with a simple nervous system but no centralized information processing, to improve the understanding of nervous system evolution. The N4 neuronal circuit in embryos develops through activity-driven self-assembly, where neurons in distinct regions increase connectivity and synchronization. Gap junctions and vesicle-mediated communication between neuronal and non-neuronal cells drive rapid assembly, with the embryo’s prospective oral region exhibiting the highest neuronal density. An artificial electrical circuit-based model demonstrates dynamic increases in synchronization over time, along with predictions for selective dynamic adaptions of connections. Environmental factors, like temperature and an absent microbiome, modify neural architecture, suggesting the existence of a certain plasticity in neural development. We propose that these fundamental features originated in the last common bilaterian ancestor, supporting the hypothesis that the basic architecture of the nervous system is universal.

## Main

Neuroscience is characterized by the inherent complexity of the nervous system. In extant bilaterian animals, various types of neurons establish thousands of synaptic connections that form numerous neural circuits. These allow receiving sensory input, signal propagation, decision-making, and motor output, thereby enabling multifaceted behaviors. The mechanisms that control the development and structure of functional neural circuits remain poorly defined, although they broadly involve neuronal connections^1^. Given the complexity of bilaterian neural circuits, it is essential to explore the development, structure and function of neural circuits in simpler, genetically accessible model systems that can also provide insights into the evolutionary history of neural circuitry.

Recently, with the advent of developmental neuroimaging and functional, quantitative approaches, the nervous system of the ancestral metazoan *Hydra* has proven to be suitable for investigating the origins and functions of neural circuits. As a member of the phylum Cnidaria, *Hydra* occupy a key phylogenetic position for examining the emergence of the nervous system (**Fig. 1a**). Cnidaria are our most distant bilaterian relatives and the molecular architecture of their nerve cells shares significant similarities with other invertebrates and also vertebrates*^2^*.

**Fig. 1:**
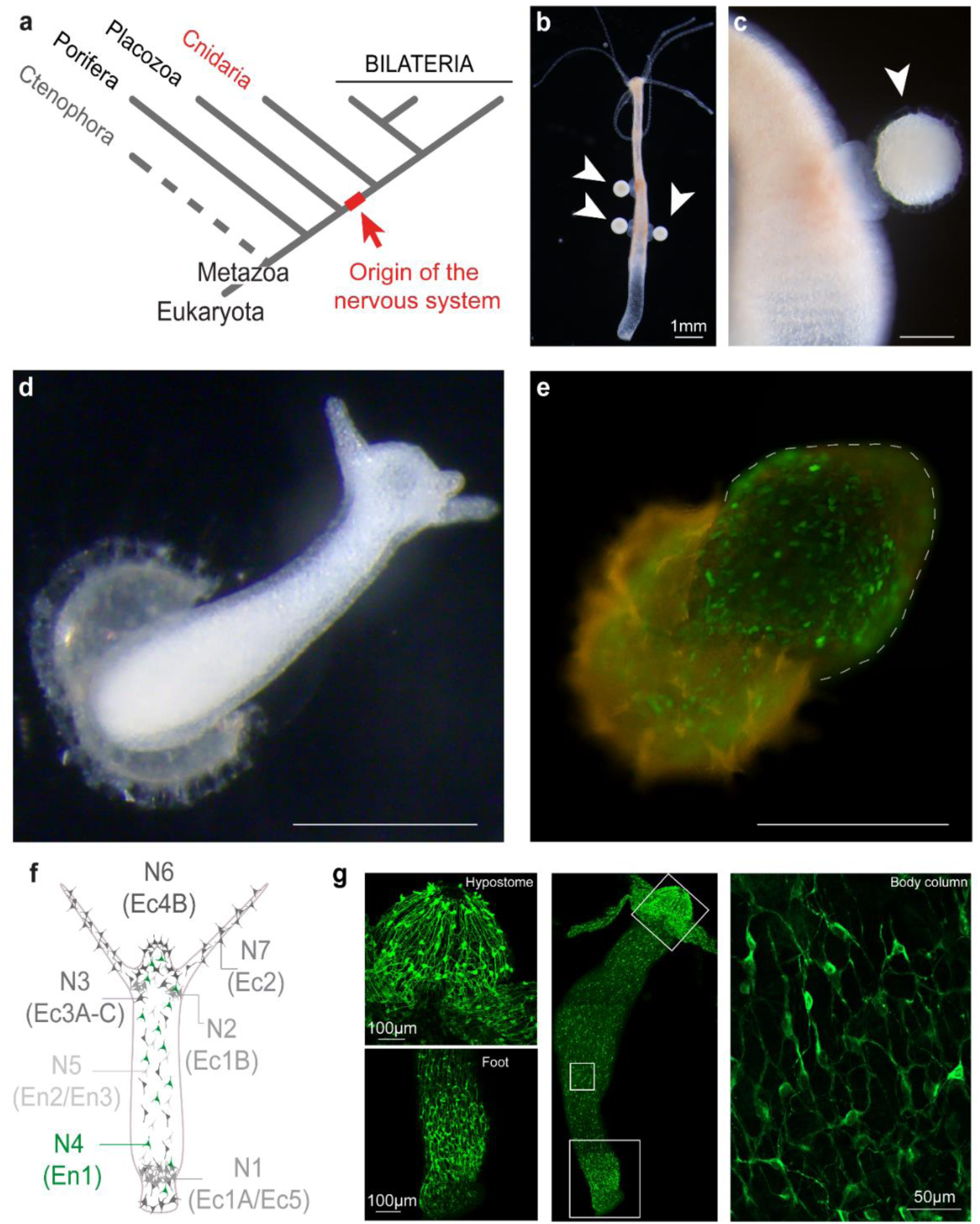
Establishing *Hydra* embryos to decode nerve net development. **a,**Phylogenetic positioning of *Hydra* with an evolutionary close standing to the origin of the nervous system. **b,c,** Female *Hydra* polyp carrying multiple fertilized embryos (white arrows) during sexual reproduction. **d,** Early hatchling, still partially covered by the cuticle, exhibiting high morphological similarity to adult polyps. **e,** Transgenic polyp^25^ in the process of hatching. Transgenic embryos provide, for the first time, insights into the development of an evolutionary ancient nerve net. **f,** Schematic overview of distinct neuronal populations categorized through RNA-sequencing. **g,** Immunohistochemical analysis of a pan-neuronal Ca^2+^ activity line (P46) generated by cross-breeding of various lines for specific neuronal populations. Close magnification on foot and hypostome (left), total view of a whole polyp (middle) and neurons forming a dense network in the body column (right).

With its diffuse nerve net consisting of around 3000-5000 neurons^3^ *Hydra* is able to display a rich repertoire of behaviors like contractions, somersaults, eating, floating and bending. The radially symmetric body is composed of two neuronal networks, located in the endo- and ectoderm, each containing distinct neuronal populations^4,5^. Both networks are separated by a structure called mesoglea without any direct bridging neuronal connections^6,7^ Neurons are located throughout all parts of the polyp’s body with a higher density in the oral and aboral regions. However, only two populations appear to encompass the entire animal, each confined to either endoderm or ectoderm. New imaging^8^ and sequencing technologies^4,5,9^, transgenic lines^10–12^ and bioinformatic pipelines allow exploration of this evolutionary ancient nervous system at single cell resolution. In particular, genetic access to distinct functional subtypes that comprise the *Hydra* nervous system have already led to remarkable new insights into the structure and function of the nerve net in adult *Hydra*. Network architecture and neurogenesis^6,7,12–14^ in adult polyps are currently the subject of intense research. Reaggregation experiments combined with Ca^2+^ imaging have proven feasible^13^ for examining a reconnecting nerve net. Recent studies could also demonstrate the neuronal influence on behaviors like feeding^12^, somersaulting^15^, phototaxis^6,16^ or detachment and even metabolic states like satiety^6^.

All these findings clearly show that the ancestral nervous system of *Hydra* is anything but primitive, instead, it comprises numerous subpopulations, each forming its own neural circuit, occupying specific localization in the body, and interacting closely with the other circuits. Here, a neural circuit is defined as a population of interconnected neurons whose loosely synchronized firing patterns carry out a specific function when activated^17,18^. Multiple neural circuits interconnect to form large-scale neural nets, and many of these nerve nets collectively constitute brains and nervous systems^19–24^.

How these neural circuits assemble *de novo* during embryogenesis in an ancestral metazoan and how they function has not yet been investigated. Here, we extend the *Hydra* research platform to include embryos and demonstrate how a neural circuit forms *de novo* in *Hydra* embryos (**Fig. 1b-e**) over developmental time. We focus on the generation of complex spatiotemporal activity patterns, as well as their functionality and synchronization level, by utilizing long-term, whole-body Ca^2+^ imaging at single-cell resolution (**Fig 1 f,g**) during late embryogenesis and early hatchlings coupled with bioinformatic analysis. To assess the variability of the neuronal architecture, embryos were exposed to different external environmental influences.

We demonstrate that the expansion of a neural circuit is facilitated by an activity-driven self-assembly and an increase in synchronization and connectivity in spatially distinct regions of the embryo. The prospective oral region of the embryo has the highest density of neurons and the strongest connections. Furthermore, an electrical circuit-based model of a self-organizing system visualizes the gradual increase in the synchronization levels over time and indicates dynamic adaptions of individual connections. In addition to high temperature, the absence of the colonizing microbiome alters the architecture and connectivity of the neural circuit, and thus indicates that even such simple and basic neural circuits possess the ability of structural modifications.

## Results

### Ultrastructural analysis of neurons in Hydra embryos at specific developmental stages

To investigate structural nerve net characteristics in late-stage embryos and early *Hydra* hatchlings, we first defined specific developmental stages based on morphological changes occurring before and after hatching (**Fig. 2a**). Here, the Bilayer stage represents the final phase of embryogenesis and the earliest stage accessible for experiments. The formation of the ectoderm is completed during this stage, but the embryos remain covered by a thick protective cuticle which was removed prior to all experiments. The Hatching stage is marked by the rupture of the cuticle, allowing the spherical embryo to elongate by incorporating surrounding medium, transitioning into a cylindrical shape (Cigar stage). The initial contractile behavior induces a significant morphological transformation, during which two small tentacles and the hypostome become visible (Hypostome stage). Within the next hours, the early hatchling exhibits a set of behaviors including periodic contractions, bending and somersaults similar to adult polyps. These behaviors are followed by the extension of the tentacle and an inducible mouth opening (Tentacle stage).

**Fig. 2:**
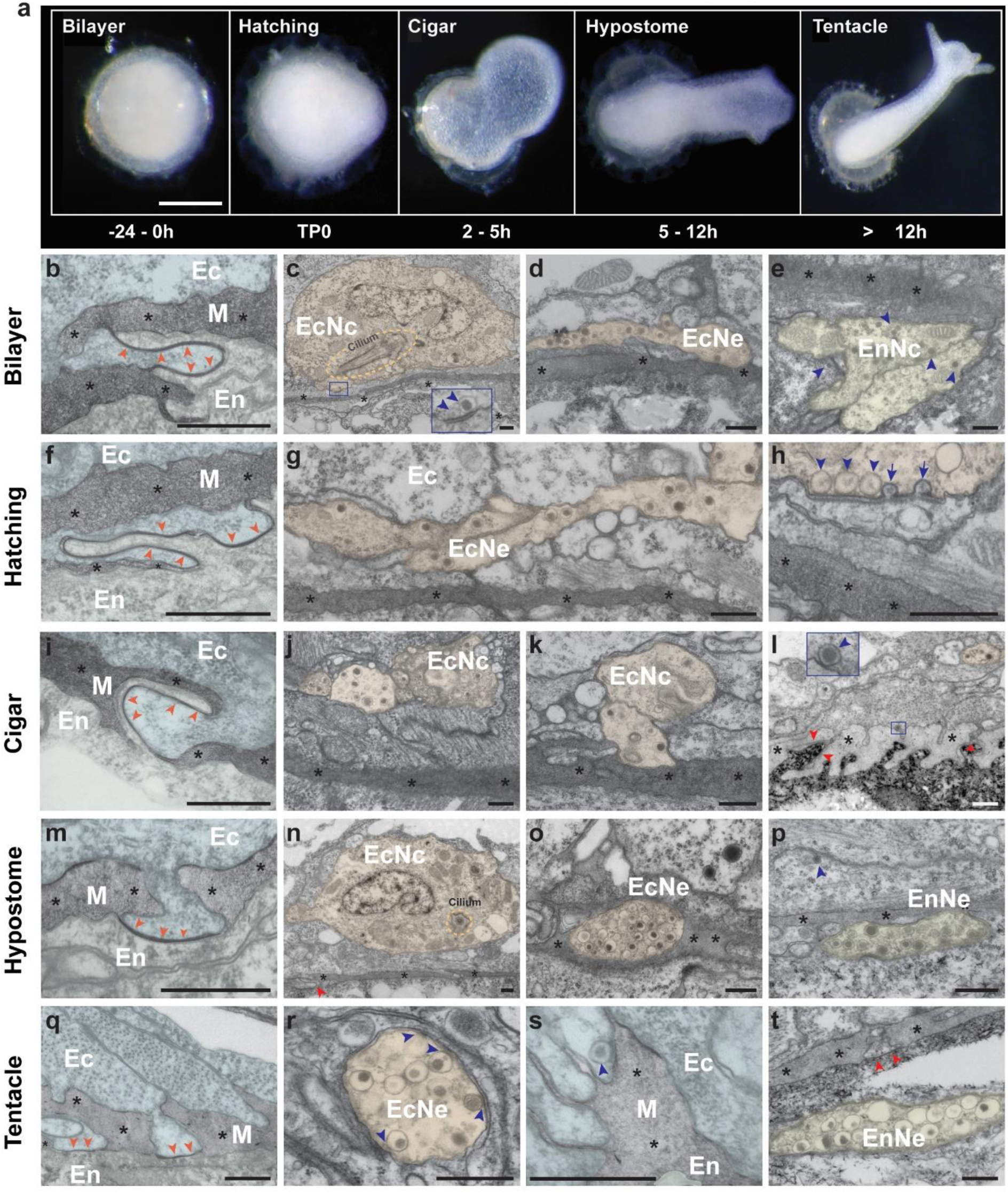
Morphology and ultrastructural analysis of early Hydra hatchlings. **a,**Sampling timepoints based on morphological changes during development of *Hydra*. In the Bilayer stage the ectodermal layer is formed, which marks the latest embryogenic stage and thereby displays the earliest accessible stage. Scale bars, 500 μm. **b-t,** Longitudinal resin sections of early hatchlings analyzed with TEM. Images are orientated to display the ectoderm above the mesoglea, below the endoderm. Structures like ectodermal and endodermal neurons and neurites characteristic for the adult nervous system were observed as early as the Bilayer stage and increased in frequency during development. Gap junctions (red arrows) between epithelial cells and electron dense vesicles (blue arrows) in nerve net and non-neuronal compartments represent a potential way to communicate between ecto- and endoderm. Scale bars, 500nm. (For more information and uncolored sections see Extended DataFig. 1-4) Ec, ectoderm; En, endoderm; M, mesoglea (*); EcNc, ectodermal neuron; EnNc, endodermal neuron; EcNe, ectodermal neurites; EnNe, endodermal neurites; red arrows, gap junctions; blue arrows, electron dense vesicles (EDV).

Following these stages, we performed transmission electron microscopy (TEM) analysis by imaging longitudinal resin sections to gain insights into the nerve net structure and the connectivity during the growth of *Hydra* hatchlings (**Fig. 2b-t**, Extended Data Fig.1b). Structures characteristic for the adult nerve net were observed as early as in the Bilayer stage and increased in frequency during development. Gap junctions (red arrowheads) were found interrupting the mesoglea and were characterized by closely apposed plasma membranes of epithelial cells separated only by a narrow electron-dense layer (**Fig.2 b,f,I,m,q**, Extended Data Fig. 2a-d). Additionally, neuron-like cells were identified based on distinguishing features such as cilia, a compact nucleus (**Fig.2 c, n**) and electron-dense vesicles (EDV) (**Fig. 2h**, Extended Data Fig2d-g). Cell bodies of neuron-like cells found in the endoderm (**Fig.2 c, n**), and neurites localized either in the ectoderm (**Fig. 2d,g,j,k**) or endoderm (**Fig. 2e,p,t**) were separated from the mesoglea by muscular processes. Occasionally, neurites were observed in contact with the mesoglea, with some exhibiting a high density of EDV (**Fig. 2d,k,o,p**). Furthermore, we identified EDV within ectodermal epithelia cells (**Fig. 2I,p,s**) docked to the plasma membrane (**Fig. 2h**; Extended Data Fig.2d-g), directly exposed to the mesoglea in the polar regions of later staged samples (**Fig. 2l,s**) but at lower abundance compared to neuronal structures. Notably, EDV appeared to be prominent in neuronal compartments during early development and were frequently observed in close proximity to the plasma membrane of cells likely to be non-neuronal (**Fig. 2c,h,r,t**), suggesting a potential role in facilitating cell-cell communication. As maturation progresses, polar regions, which indicates the early formation of a head or foot could be identified by numerous extensions of ecto- and endodermal cells crossing the mesoglea in polyps of the Cigar (Fig. 2l) or Tentacle stage (**Fig. 2s**). Additionally, the extensions of the ecto- and endodermal muscular processes appeared to interact within the mesoglea via gap junctions (Extended Data Fig.3). Analyzing the later staged sections, our analysis revealed signs of neuronal maturation, supported by maceration (Extended Data Fig.1 c-e). This was evidenced by an increase in heterochromatin (**Fig.2 c,n**) and the accumulation of electron-dense vesicles in neurites (**Fig.2 o,p,r,t**) indicating a transition to a more differentiated neuronal state.

Collectively, these results provide novel insights into the connectivity and potential signaling mechanisms in *Hydra* embryos. The rapid assembly of a functional nerve net capable of facilitating complex behaviors may be driven by gap junctions as well as vesicle-mediated communication between nerve cells, transmitting signal molecules throughout the mesoglea and non-neuronal cells, indicating their pivotal role in nervous system development. While, in agreement with previous observations^6^, we found no evidence for direct connections between the endodermal and ectodermal nerve nets, epithelial cells were observed to remain in close contact across the mesoglea as described previously^6^. Ectodermal and endodermal epithelial cell processes can enter and even traverse the mesoglea. Furthermore, nerve cells within the ecto- and endodermal layer are in contact with mesoglea and epithelial cells as well as other neurons within each layer.

### Topology of neurons in developing embryos

To measure, further characterize, and visualize the early nerve net structure based on different stages (**Fig.2 a**) we stained distinct neuronal populations by using a combination of the specific neuropeptides GLWamide (exclusive to population N2, N3, N4) and RFamide (exclusive to population N1, N6, N7) in wild-type samples. Our findings indicate a fully developed and interconnected nerve net throughout the hatchlings during the early developmental stages as well as a distinct oral and aboral patterning (**Fig.3 b,c**, Extended Data Fig.5 a,b). The latter was achieved by visualizing neuronal populations that are exclusively expressed in the hypostome and foot region of adult polyps^4,6,26^.

**Fig. 3:**
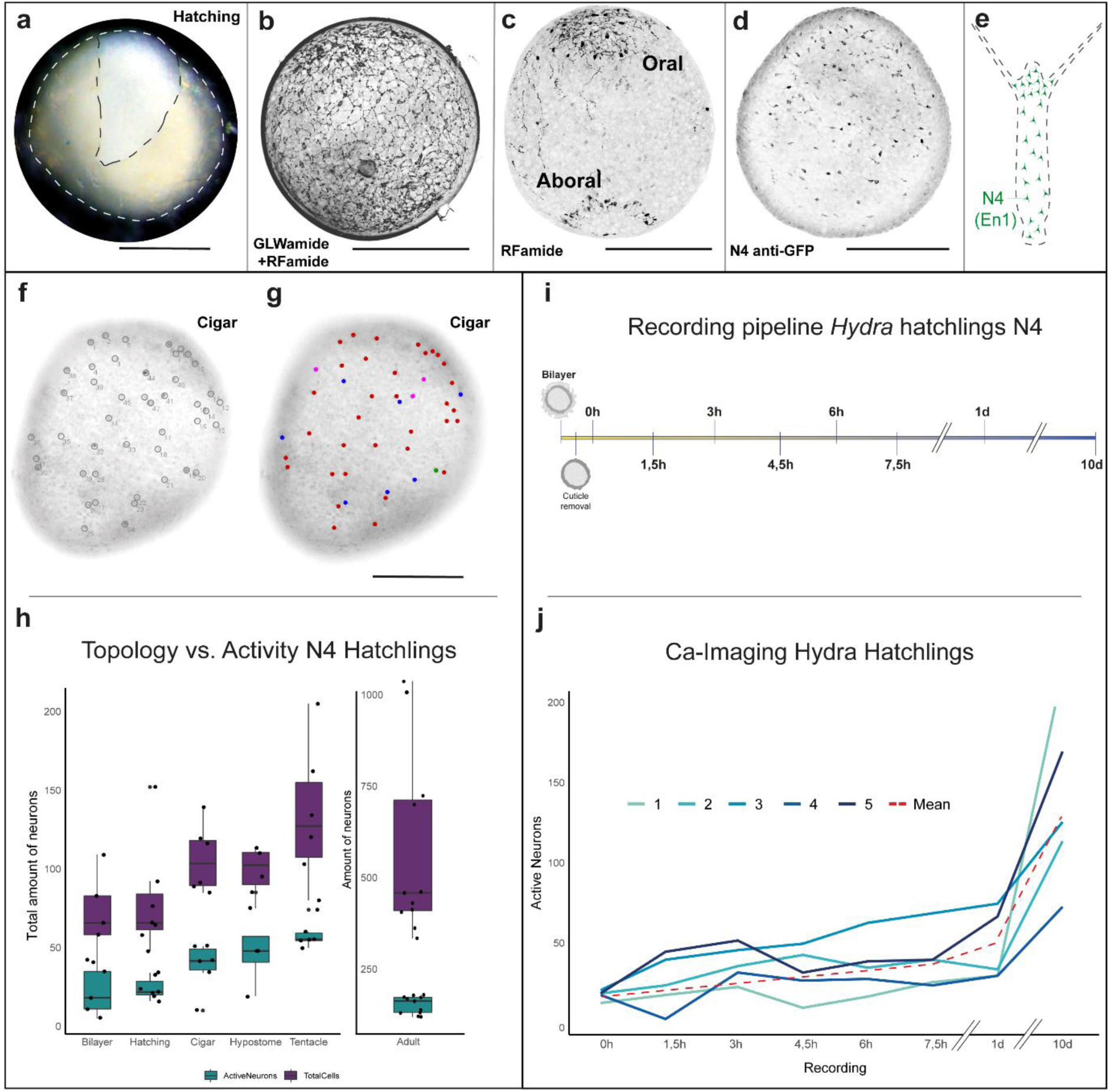
Encoding nerve net topology and establishment of transgenic embryos. **a,**Rupture of the cuticle (dashed black line) in the Hatching stage. **b,** Staining against two specific neuropeptides (GLWamide, RFamide), expressed in *Hydra* neurons, delivered first insights into the whole nervous system during Hatching stage in wild-type polyps. **c,** RFamide staining revealed an early patterning already during Hatching stage by staining neuronal populations specific for the future head and foot. **d,** GFP immunolabelling revealed a successful initial implementation of transgenic *Hydra* embryos for calcium imaging (GCaMP6s) of the endodermal N4 population. **e,** Schematic distribution of the endodermal N4 network in adult polyps illustrating the highest density in the hypostome area and no expression in the tentacles. **f,** Example of fixed hatchling in Cigar stage showing labelled N4 neurons with anti-GFP antibody. **g,** Different-sized and color-coded neuronal ensembles analyzed by Ca^2+^ activity before fixation. Red dots mark the dominant N4 community. **h,** Quantification and comparison of active and stained N4 neurons at different developmental stages revealed lower numbers of spiking neurons compared to those expressing the N4-specific transcript, targeted by antibody staining. The number of neurons counted for the visual field in both treatments, equivalent to one half of the samples display an overall increase during development. (for topology n = 6-11; for activity n = 4-11) **i,** Recording pipeline for long-term analysis of single *Hydra* samples starting in the de-cuticulized Bilayer stage. Self-made observation chambers ensured a permanent screening throughout the whole experiment until 7.5h after start. Recordings were made for 30min with 1h break. **j,** Ca^2+^ active N4 neurons analyzed for each recording in 5 individuals within 2000 frames of 30min recordings. The mean for all measurements displays an overall increase of neuronal activity (dashed red line). Scale bars 500μm.

### Ca^2+^ imaging reveals the location and number of active N4 neurons in the embryo

To gain a more profound understanding of the network formation in an emerging nervous system we utilized transgenic embryos, focusing on the endodermal population N4 (**Fig. 3d,f,g**). Initially, we analyzed Ca^2+^ activity for the previously described developmental stages. In a subsequent step, we performed antibody staining of the same samples against GFP as a compound of the GCaMP6S cassette to compare the spatial topology of neurons and the activity of the N4-specific population. Analysis of the Ca^2+^ signals revealed the presence of neuronal ensembles of varying sizes throughout all regions of our samples (**Fig. 3f,g**, Extended Data Vid1-4). Comparing the results, we observed an increase of both total and Ca^2+^ active neurons (**Fig. 3h**). Notably, we observed a lower number of active neurons during Ca^2+^ imaging compared to the total number of N4-positive cells identified via Immunohistochemistry. Specifically, in early stages, we identified an active-to-total cell ratio of 1:3, which shifted to a ratio of 1:2 in later stages, before returning to the initial proportion seen in adult polyps, similar to the Bilayer stage. This finding indicates a declining pool of potentially immature neurons being integrated into an expanding Ca^2+^ active network (**Fig. 3h**, Extended Data Fig. 5c). Since this approach does not allow for sustained observation of the evolving network structure, we developed a pipeline for a continuous recording that maintains focus on the same section of individual polyps, thereby avoiding deviations in the collected data (**Fig. 3i**). The resulting single-cell measurements demonstrated an increase in the number of active neurons over a 10-day period, specifically focusing on the first 7.5 hours of development in our analyzed samples (**Fig. 3j**).

### Ca^2+^ activity drives the formation and synchronization of the N4 neural circuit

To investigate the assembly of the evolving N4 neural circuit, we monitored neuronal activity by tracing all detectable neurons on a single-cell basis for 2000 frames at 25fps during the initial hours after cuticle removal in the Bilayer stage, followed by a detailed analysis of 1000 frames at 1 and 10 days after extracting the embryos from the cuticle. Overall, we analyzed between 12 individual neurons in early-staged embryos and 143 neurons in adult polyps. Neuronal activity was quantified by assessing the intensity of each individual neuron and the resulting data were compared across timepoints to evaluate synchronization levels (**Fig. 4a**). Our results suggest that, rather than simply an increase in the number of neurons over time, there is an enhanced synchronization among Ca^2+^ active neurons, culminating in predominantly simultaneous activity after 1 and 10 days. Interestingly, we observed an increase in neuronal spiking rates between 0- and 7.5-hours post-cuticle removal, whereas we exhibited an enhanced synchronization level by detecting a decrease in calcium spike frequency after one day (Extended Data Fig. 6a). This suggests a high spiking activity as crucial for the early stages of nerve net assembly and indicating a finalized network construction after 1 day. Collectively, these findings suggest a rapid incorporation of new neurons and a corresponding increase in connectivity during early developmental stages, promoting the expansion of the network. Furthermore, our recordings revealed variable numbers of Ca^2+^ active neuronal ensembles and different levels of connectivity in distinct regions of the hatchlings (**Fig. 3g**, Supplementary Video 1-4), implying the absence of a chemical or structural pre-patterning and instead supporting an activity-driven assembly.

**Fig. 4:**
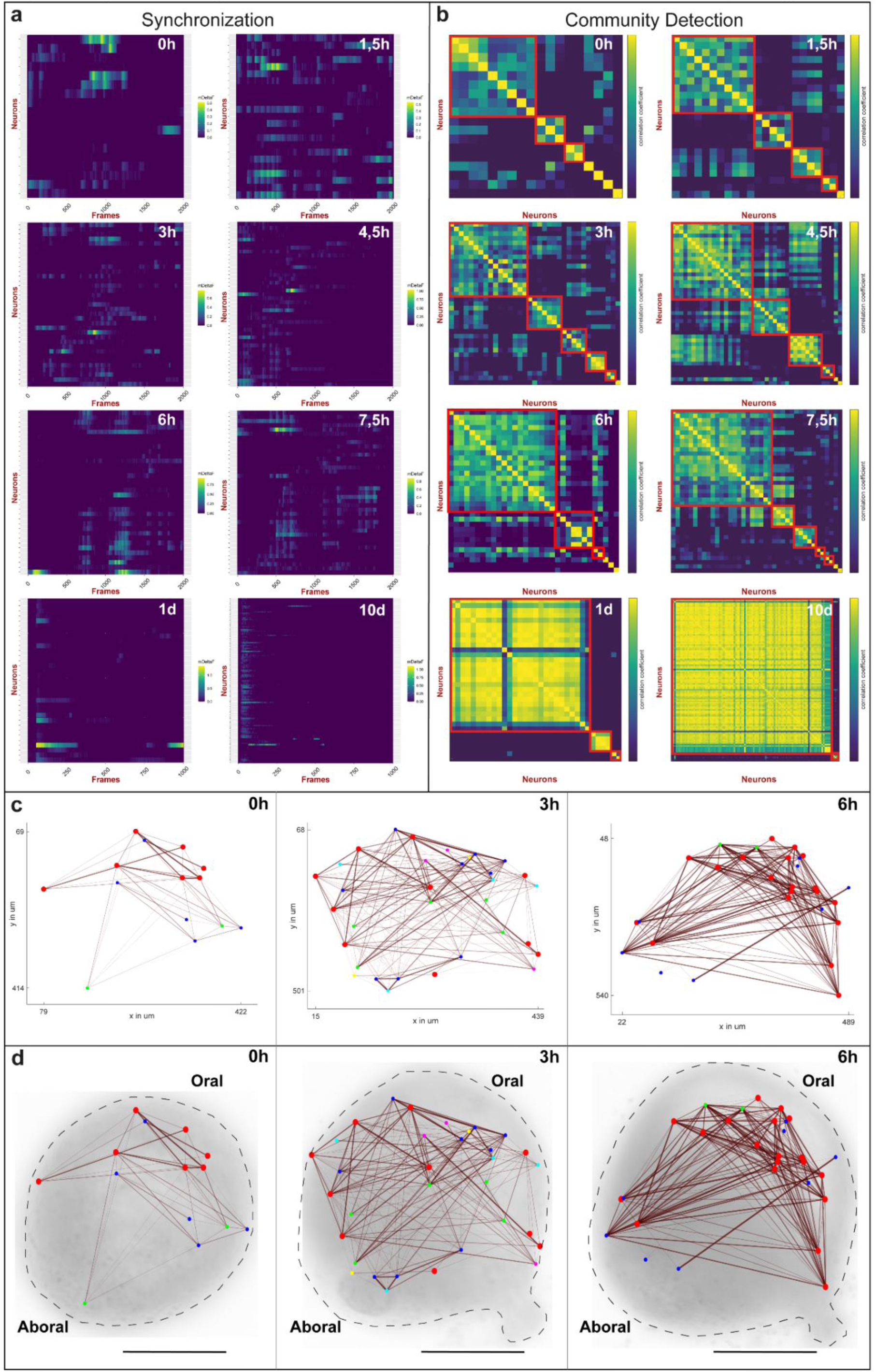
Single-cell analysis of neuronal activity unmask nerve net assembly in *Hydra*. **a,**Single-cell analysis of neuronal activity during recordings indicate an increase of synchronization level and a high number of spikes in early developmental stages (0 – 7,5h) compared to later stages (1d, 10d). Each line on the y-axis corresponds to a single detected neuron, x-axis shows the spike intensity (ΔF/F) for each frame (25fps). **b,** Utilizing the data Ca^2+^ measurements presented in **a,** a community detection for each recording to follow the network assembly was performed Detected communities are highlighted by red squares identified with the Leiden algorithm using the correlation coefficient to visualize the rapid ongoing network assembly in *Hydra*. Throughout the analysis, the dominant community increased in size, along with an enhancement in the correlation coefficient, while smaller communities and single neurons remained detectable in all time points. In the late stages (1d, 10d), the consistently high correlation coefficient indicated the completion of the network assembly. **c,d,** The network assembly visualized by using the positional identity of detected neurons. Colored dots represent distinct neuronal ensembles based on the correlation coefficient depicted in **b** (red = dominant community). Connections between neurons are shown by lines, the thickness reflecting connection strength. Together, these data provide evidence for a rapid assembly of the N4 neural circuit, a rise in strongly connected neurons within the dominant ensemble and an aggregation of neurons in the prospective oral region of the hatchling.

To further examine the network construction, we performed a community detection based on the Leiden algorithm^27^ (**Fig. 4b**). Our analysis identified neuronal ensembles of varying sizes and numbers, as well as isolated neurons, particularly during early stages of development (Extended Data Fig.6). Notably, the correlation coefficient increased over time until mostly all detected neurons exhibited a strong functional connectivity by belonging to the spanning N4 network after 1 or 10 days. A conspicuous feature in our analysis was the emergence of a dominant community during the study period. This neuronal ensemble, constituting approximately 50% of the traced neurons, increased in size over time. We hypothesize that this large neuronal ensemble functions as a pacemaker-like structure, overriding smaller ensembles due to its activity and spiking intensity. It appears to incorporate smaller, already-connected and synchronized ensembles, thereby facilitating rapid circuit expansion.

To achieve a deeper understanding of the time-dependent nerve net assembly in the N4 network, we employed a 2D network analysis that uses the positional identity to reconstruct the geometric network structure. We could, thereby, gain further insights into modularity, dynamic adaptions, connectivity and integration of isolated neurons and ensembles. Building on previous findings (**Fig. 4a,b**), we observed an activity-driven self-assembly of the nerve net during recordings (**Fig. 4c**, Extended Data Fig. 7). Our analysis revealed an increase in connectivity within specific ensembles but likewise between diverse neuronal clusters throughout our analysis. In alignment with our earlier results, we detected growth in the dominant neuronal community through the incorporation of additional ensembles particularly in the upper region of the nerve net. Furthermore, we observed not only an increase in connections but also selective strengthening of specific connections, especially within the dominant community (marked by red dots) over a remarkably short period of time. Interestingly, smaller neuronal ensembles appeared to disappear, after getting locally integrated and synchronized within the main network as the larger ensemble expanded and matured, suggesting a dynamic reorganization and growth of the nerve net Intriguingly, the highest density of neurons and the establishment of the strongest connections were concentrated in the oral region of the recorded hatchlings (**Fig. 4d**). This suggests that the origin of the N4 circuit assembly is located in the future hypostome, where the N4 concentration reaches the highest density also in adult polyps^6,12^).

### An artificial electrical circuit-based model simulates neural circuit formation and predicts dynamic adaptions

To gain an even deeper insight into the formation of the N4 neural circuit, we have developed a first “observer” model using an electrical circuit-based approach. As illustrated in **Fig.5a**, although the model operates over a shorter time span compared to our *in vivo* studies, yet specific observation points align closely with those made in the *Hydra* embryos. The electrical circuit-based model allows us to continuously display calcium activity in real-time, here derived directly from electric signals (**Fig. 5b**). Applying the previously used community detection algorithm and separating our examinations into 20-second windows (**Fig. 5c,d**), we came to results that were remarkably similar to our earlier *in vivo* observations (see **Fig. 4b**). Results achieved by a circuit model with identical parameters representing neurons (**Fig. 5c**) especially reflect the rise in the synchronization levels during development, although a single large network forms directly. In contrast, results for a deeper modeling with varying parameters that represent different membrane capacitances, leading to diverse neuronal firing rates (**Fig. 5d**), also show the initial formation of smaller sub-networks before they merge into one global network. This highlights an important insight: while parameter variations in circuit design are typically considered non-ideal, they are essential for the emergence of distinct neuronal sub-networks as observed in biological systems. Thus, non-idealities in electrical circuits and oscillators such as parameter variations may in fact be advantageous, enabling the formation of distinct sub-networks with different functionalities, rather than a single network serving a singular function.

**Fig. 5:**
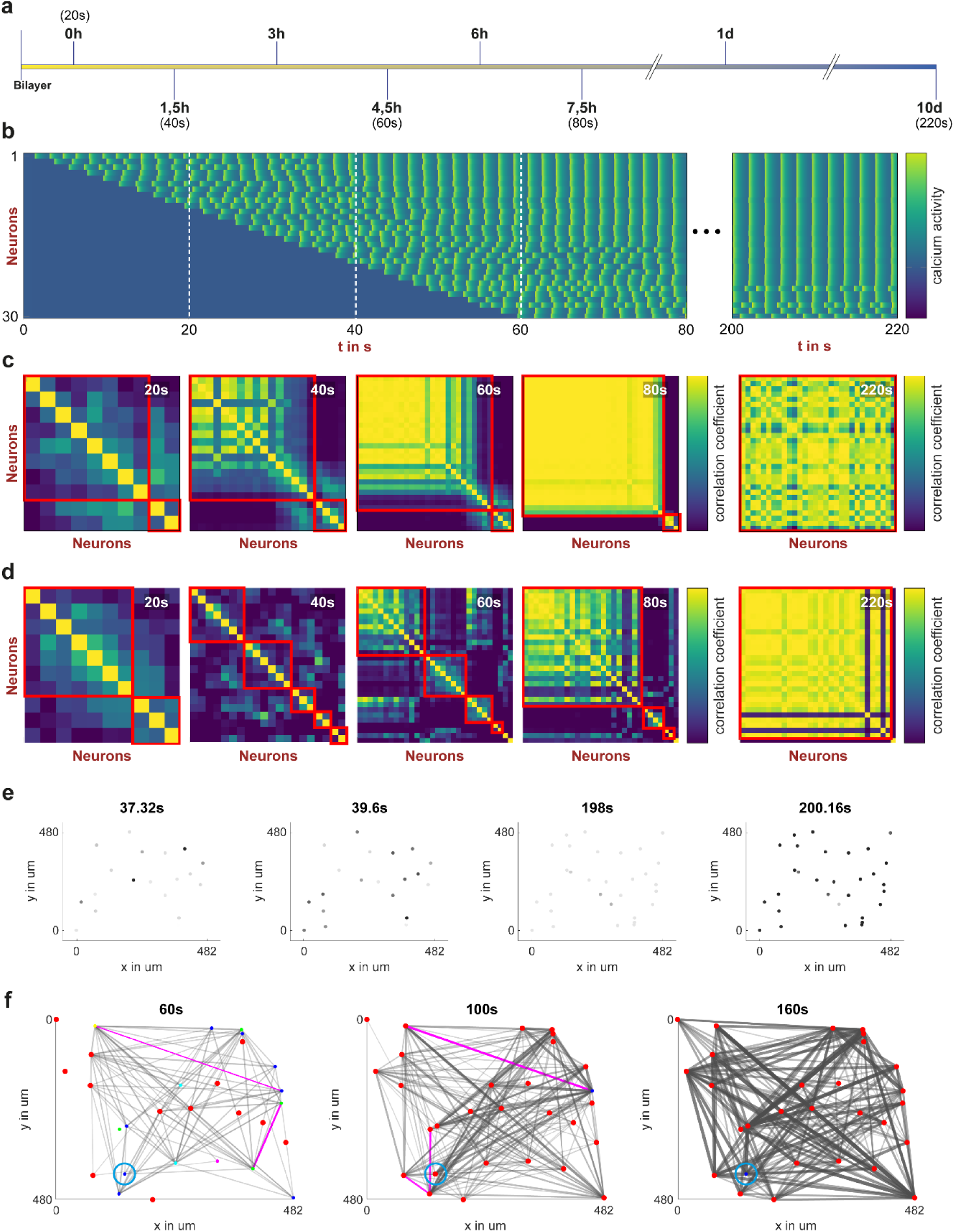
Artificial circuit simulations enable N4 network-specific predictions. **a,**Adaption of the recording pipeline for *Hydra* hatchlings and the simulation. **b,** Simulated calcium activity. Simulated neurons were integrated at 2-seconds intervals until reaching a total number of 30. **c,** Detected communities in a simulated nerve net with identical, ideal neuronal parameters, highlighted by red squares. **d,** Detected communities for the simulated activity of **a** using varying spike frequencies based on different membrane capacitances. **e,** Simulated brightness levels of **a** exhibit an increase in synchronization. **f,** Real connections within the simulated nerve net shown in **a**. Thickness of lines represents the connection strength between simulated neurons. Structural changes, indicative for dynamic adaptions, are demonstrated by assembly and disassembly of neuronal connections (pink lines). Additionally, network dynamics are illustrated by the integration and disintegration of individual neurons (blue circle).

Additional advantages of the circuit as an observer model include the availability of position data for all neurons in simulations and the capability to generate artificial calcium-imaging videos (**Fig. 5e**, Extended Data Fig.8, Vid2). This allows us to visualize the gradual increase in the synchronization levels over time. Specifically, at 40 seconds, different neurons begin firing at two consecutive time points, while at 200 seconds, nearly all neurons fire synchronously.

The circuit’s observer capabilities are further demonstrated by its ability to simulate evolving connections, offering direct access to real-time network changes (**Fig. 5f**). In contrast to network graphs derived from experimental data, which rely on estimations of correlation analyses, the circuit model captures the actual dynamics of connection formation. This allows us to investigate and track gradual changes in the network structure that may not be fully captured in the recorded data. Notably, we observed integration and disintegration of neurons into the dominant community and dynamic adaptions of individual connections, indicating a dynamic network even though the overall structure remains similar. These dynamic adjustments and the removal of redundant or otherwise unnecessary connections might explain a slightly decreasing number of neurons observed in our previous measurements (see **Fig. 3j**), as reassembly of connections may occur, but has not been observed in detail yet.

### Temperature and absence of microbiota impact formation and architecture of the N4 circuit

To investigate whether external factors in the immediate environment of embryos affect the structure of the N4 neural circuit, we examined two external factors: temperature and potential signals from the colonizing microbiome. To ensure normal embryonic development, the cuticle was retained intact until the embryos reached the Bilayer stage. To first investigate the influence of temperature, the embryos were transferred from the standard temperature (18°C) to either warm (23°C) or cold (13°C) culture medium immediately after removal of the cuticle. To analyze the Ca^2+^ activity we used our previously implemented pipeline (see **Fig. 3i**) focusing on early stages (**Fig. 6a,b**, Extended Data Fig.9,10). Interestingly, we observed significant deviations from our measurements under standard conditions (18°C, see **Fig. 4**) for both the average community size and the spiking frequency. Hatchlings exposed to cold temperature (13°C) exhibited significantly increased synchronization levels and enhanced neuronal connectivity within the dominant community compared to the control group (**Fig. 6a,c**), indicating that cold temperature promotes a rapid neuronal network assembly. In stark contrast, hatchlings recorded at high temperatures (23°C) exhibited reduced synchronization levels. The community detection in embryos at high temperatures revealed that the dominant neuronal ensemble was smaller in size compared to the total number of detected neurons (**Fig. 6.b,c**). Remarkably, these structural and functional changes in the N4 neural circuit directly influenced the behavior of the hatchlings, as 40% of hatchlings maintained at 23°C were unable to engulf prey and did not survive to the end of the 10-day experiment (**Fig. 6c**, BarChart). This is very strongly reminiscent of the behavioral defects in our previous study, where we ablated the N4 population in adult polyps^6^. In addition, Ca^2+^ signals recorded in N4 neurons of embryos kept at warm temperatures indicated an increase in spiking frequency. Incorporating the temperature-driven alterations in the spiking frequency of the N4 neurons into our electrical circuit-based model, the model also predicts that the dominant neuronal ensemble is smaller in size compared to the total number of detected neurons (Extended Data Fig. 8c,d). The model can thus effectively emulate biological processes. Taken together, the data clearly indicate that embryos at high temperature face difficulties in assembling a functional N4 neural circuit, whereas cold temperatures indicate a beneficial environment for the nerve net assembly.

**Fig. 6:**
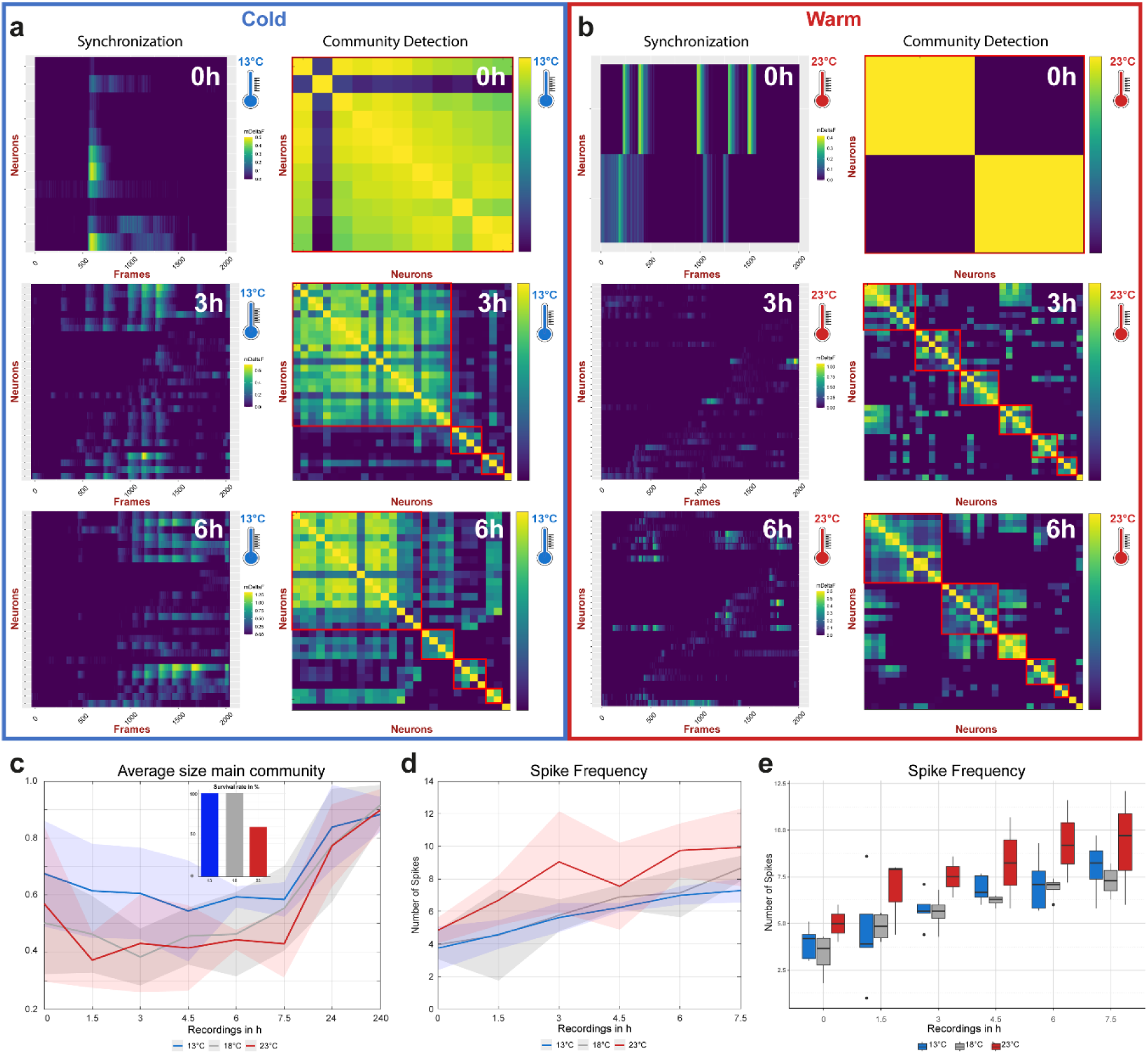
Temperature-dependent network assembly in early stages. **a,b,** Single-cell analysis of neurons recorded under cold and warm conditions starting in the Bilayer stage. Three representative early-staged recordings during development are shown. Significant differences were observed between both treatments in terms of synchronization and community detection. Reduced temperature conditions were found to be beneficial for synchronization levels during nerve net assembly, as well as convenient for the rapid development of a dominant ensemble. In contrast, synchronization level and community size in **b** appear to be lower for those hatchlings recorded in a warm environment. **c,** Specific observations of the dominant community across all treatments (13°C – blue, 18°C – grey, 23°C – red) indicate a higher proportion of neurons connected within the main community: This suggests an increased assembly during early development under cold conditions, but did not display an advantage after 10d. In comparison, hatchlings exposed to normal and warm temperatures showed similar community sizes, except at 7,5h, where normal conditions exhibited a higher number of assembled neurons (shaded area represents the mean ± SD). Additionally, data presented in the Barchart, show a decreased survival rate of polyps kept at warm temperatures to engulf prey (n = 15). **d,e,** Spike frequency analysis revealed an increased number of signals in hatchling observed at 23°C. (shaded area represents the mean ± SD).

The second environmental factor we investigated was the influence of the microbiome on the number of RFamide-positive neurons in the oral area of early hatchlings. Based on previous findings in adult *Hydra* showing the manifold influence of the microbiome on development, fitness, and also on the nervous system and behavior^12,28–31^, we hypothesized that the absence of the microbiome may affect the *de novo* network assembly. Here, the role of the microbiota was uncovered by the absence of the natural microbiota after cuticle formation. For this, germ-free (gf) animals were compared with their wild-type (wt) counterparts and recolonized with a full complement of the microbiota (rc). Intriguingly, embryos cultured under bacteria-free condition displayed a decreased hatching rate and a drastically reduced number of RFamide positive neurons in the future hypostome (**Fig 7a,b**). Exposing gf embryos to bacteria from wt *Hydra* could not significantly reverse this effect but indicated a higher number of neurons throughout all developmental stages compared to the germ-free conditions (**Fig 7b**). To further investigate the effects of the microbiome, we analyzed the number of neurites in neurons across our treatment groups. Remarkably, we found significant differences considering the germ-free conditions in the number of both neurons and their corresponding neurites, indicating fewer connections and a lack of maturation during development (**Fig 7c-e**). Interestingly, the recolonized samples produced an unusually large number of up to 5 neurites (**Fig 7i**), suggesting a compensation in low numbers of neurons by establishing more neurite connections to neighboring cells. Using Immunohistochemistry for RFamide in germ-free and recolonized hatchlings, it became clear that the oral nerve net is not properly formed in the absence of microbes (**Fig 7 j-l**). Neurons in embryos cultured under germ-free conditions (**Fig 7 k**) are lower in number and appear to lack contact with other neuronal cells compared to the wild-type control probably also because of the lower number of neurites. These finding suggests a pivotal role of the microbiome during neurogenesis and neural circuit formation as indicated by decreased network connectivity and stability.

**Fig.7:**
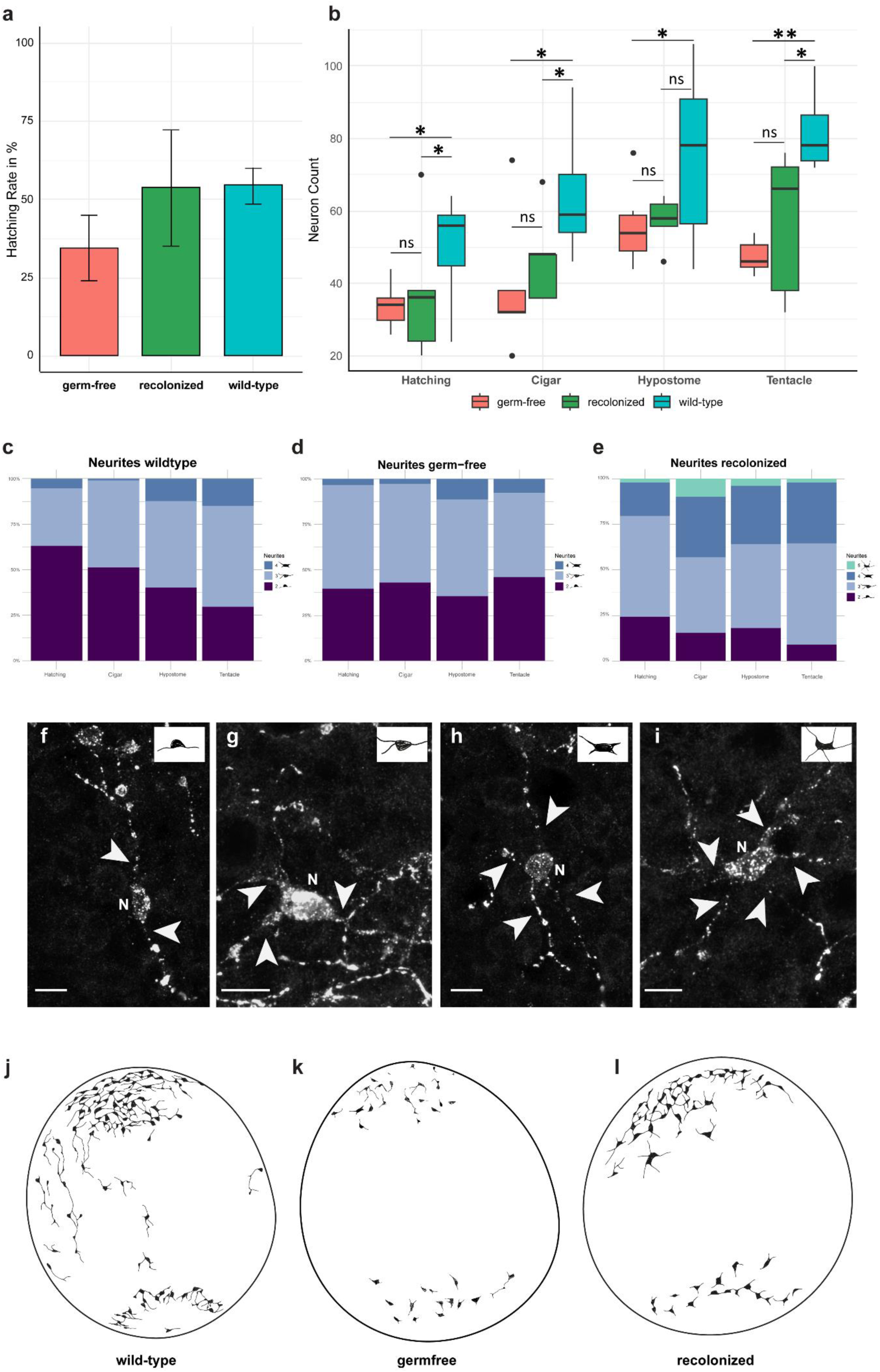
Microbial impact on the nerve net plasticity. **a,**The absence of microbes exhibited a reduction in the hatching rate of germ-free embryos compared to the wild-type. However, exposing germ-free embryos to the natural microbiome could restore the hatching rate. Bars represent the mean± sd. (wild-type: n = 3242; germ-free: n = 209; recolonized: n = 107) **b,** Neuron counts in the subsequent head, as assessed by RFamide staining, revealed a significant reduction in neuronal density compared to the wild-type control. Recolonized embryos displayed no statistically significant difference in neuron counts, but consistently showed a slight increase across all developmental stages. (statistics: ANOVA, Tukey test, Kruskal-Wallis, Dunn test; p-values: ns: ≥ 0,05; *: < 0,05; **: < 0,01) **c-e,** Analysis of neuronal branches in wildtype, germ-free and recolonized hatchlings revealed distinct variations across all treatments. In wild-type samples, the number of neurites increased as development progressed. In contrast, germ-free hatchlings maintained a stable distribution, with neurons primarily showing 2, 3 and 4 branches. Recolonized hatchlings, however, displayed a higher proportion of neurons with 3 and 4 neurites even in the early developmental stages. Notably, neurons with 5 branches were absent in both wildtype and germ-free hatchlings, appearing exclusively in recolonized individuals (wild-type: n = 5-14; germ-free: n = 6-8; recolonized: n = 5). **f-I,** Immunohistochemistry for RFamide in germ-free and recolonized hatchlings. Arrowheads indicate the neurites. N, Neuron. Scale bar 10µm. **j-l,** Schematic representations of immunohistochemistry against RFamide for wild-type, germ-free and recolonized samples in the Hatching stage (Extended Data Fig.11).

## Discussion

Since Cajaĺs observations over a century ago that neural circuits consist of neurons^32^, neural circuits have fascinated neuroscientists. The role of neural circuits of the central nervous system is well documented as regulatory pathways for feeling, motion control, learning, and memory, and their dysfunction is related to various neurodegenerative diseases^33^. Those studies, however, have not revealed the mechanisms behind the first emergence of neural circuits at the early stages of animal evolution. Our objective was to uncover how neurons in one of the oldest phylogenetic nervous systems learned to assemble into functional neural circuits and to question whether external environmental influences affect the phenotypes of developing neurons and the architecture of the entire neural circuit. We focused our work in the ancestral metazoan *Hydra* on the embryonic assembly of the N4-specific neural circuit, as the endodermal neuronal subpopulation N4 in the adult body is found throughout the animal and its presence is a pre-requisite for the eating behavior^12^. Particularly, the N4 population plays a crucial role in prey engulfment following mouth opening and the spit-out of the digestive pellet, as demonstrated in previous ablation experiments^6^. We discovered that neurons are present before they are functionally required and that the information of the neural circuit unfolds over time as a self-assembling system, driven and synchronized by Ca^2+^ signals as well as an increase of neurons. Ca imaging demonstrated that there is a steady increase in active neurons and that the formation of the neural circuit is a dynamic process. This neural circuit assembly enables modifications in connectivity patterns potentially through dynamic adaptions, and allows an adjustment of functional circuit properties in response to changing external environments.

The neural communication pathways in the embryo mainly involve gap junctions as well as vesicle-mediated communication between nerve cells, interaction with non-neuronal cells and also the involvement of the mesoglea layer (**Fig 2**). Although the number of neurons and neural collaborators and thus the possibility of interactions increases considerably in the late embryo (**Fig 3**), there appears to be no direct communication between neurons in the ectoderm and endoderm in the examined developmental stages (**Fig 2**) or, as described previously, in the adult polyps^6,7^ Although adult *Hydra* polyps are often referred to as eternal embryos^34^, investigating embryos and early-staged hatchlings offers unique insights into nerve net construction and architecture. Unlike previous experiments working on the re-assembly of the *Hydra* nerve net^13^, this study focuses on the *de novo* assembly of a distinct neural circuit from a baseline state, without the use of previously disrupted neuronal structures.

The finding that ancestral neural circuits do not start out as a randomly connected network, but form and grow by self-assembly in a dynamic process in which stability and precision are achieved through mechanisms involving a significant degree of structural and functional modulation (pruning, parameter variations, environmental modifications) may have broader implications relevant to the design of novel artificial neural networks^35,36^. Our electrical circuit-based model allows the translation of biological insights into the domain of electrical engineering and may not only facilitate the design of novel circuits but also enable the prediction of nerve net characteristics that may remain undetected in biological systems. Furthermore, the ultrastructural characterization of the neural network in the developing *Hydra* embryos revealed structural components necessary for maturation by activity-driven self-assembly, particularly gap junctions between the epithelial cells of the ecto- and endoderm, as well as the presence of EDV. The bidirectional exchange also offers to be enlarged to the implementation and progression of more complex artificial circuits simulating environmental influences by considering spatially expanding neurites based on growth or substance concentrations^37^.

While the self-assembly process leading to a functional neural circuit with inherent modifications is fascinating, we also identified an important environmental impact. Both temperature (**Fig 6**) and the colonizing microbiome (**Fig. 7**) modulate neural circuit formation and connectivity.

Numerous studies explore the effect of temperature on phenotypic differences and the impact on developmental processes^38,39^. The mechanisms by which temperature affects the formation of *Hydra’s* nervous system remain unclear. Whether the effect results from a simple alteration in chemical reaction norms or if temperature is actively sensed, leading to adjustments in developmental programs, remains a subject of debate. Temperature changes could also induce physiological effects on neurons, including modifications in membrane properties and ion pump dynamics, or affect biochemical signaling. Our findings that activity of N4 neurons is directly affected by a temperature change, encourage us to consider whether components of *Hydra’s* nervous system, neuronal or non-neuronal, can also perceive and process these signals.

Over the last two decades, *Hydra* was also developed as an excellent exploratory model organism for studying host-microbe interactions^12,40–43^. Here, we similarly observed that the absence of a microbiome results in impaired neural circuit formation, indicating that successful development and assembly depend on the continuous presence of a co-evolved microbiome. These observations support and extend a previous study in mice, which demonstrated that embryos developed a defective nervous system in the absence of maternal microbial signals^44^. Thus, the influence of the microbiome on processes of nervous system formation appears to be an ancient phenomenon that has existed since the beginning of animal phylogeny. Furthermore, while antibiotic treatment may directly affect nervous system formation, the microbiome likely plays a crucial role in mitigating additional impacts from opportunistic microorganisms such as fungi or archaea, by maintaining microbial balance. The ability to compensate neuronal deficiency in early *Hydra* hatchlings by increasing the number of connections in existing neurons by restoring the microbiome provides direct evidence of the microbiome’s importance. The microbial factors and signals involved in *Hydra’s* neural circuit assembly remain to be clarified in future studies. Determining the environmental factors that influence neural circuit assembly and stability is essential for understanding the maintenance of nerve nets and their function in the face of global environmental changes, such as increasing temperatures and loss of microbiota. Furthermore, previous studies have shown that external factors influence neural circuits during critical phases of development in organisms such as *Drosophila* or vertebrates^45^, contributing to neurological diseases^46,47^.

Across the animal kingdom, neuronal circuit formation seems to follow a universal principle long before any centralization of information processing. To ensure rapid circuit formation *Hydra* embryos and hatchlings exhibit an increased and irregular spiking activity, supporting the implementation of early behaviors. In contrast, adult polyps with full behavioral repertoire display regular spiking patterns^6^. Similarly, in *Drosophila,* locomotor behavior begins 7h before hatching, with episodic activity as the first behavior emerging 3.5h later, driven by neural activity^48^ (mod. from Kohsaka *et al.* 2012^49^). In zebrafish, spontaneous coiling, resulting from the development of early spinal circuits, appears as early as 17h after fertilization^50^ (graphic by Zoe Zorn).

Notably, similar rapid developmental processes occur in more complex animals, suggesting universal principles governing neuronal connections and nerve net assembly (“Neurons that wire together, fire together”)^23^ or postulated in the Hebbian theory^19^. Here, our results demonstrate rapid neuronal circuit development (**Fig. 4**) to achieve a functional N4 assembly. This rapid development of neural connections has also been reported in the initiation of neural connections in the spinal cord of zebrafish, which implement basic reflexes^50^, as well as in the development of the locomotive behavior in *Drosophila* larvae inducing episodic activity^49^ (**Fig. 8**). Taken together, the basic architecture of the nervous system appears to be universal and valid not only for *Hydra*, but also for vertebrates including human beings. *“We emphasize”,* at least according to Maturana and Varela^51^, “*the basic organization of the immensely complicated human nervous system follows essentially the same logic as in the humble hydra”*.

**Fig.8:**
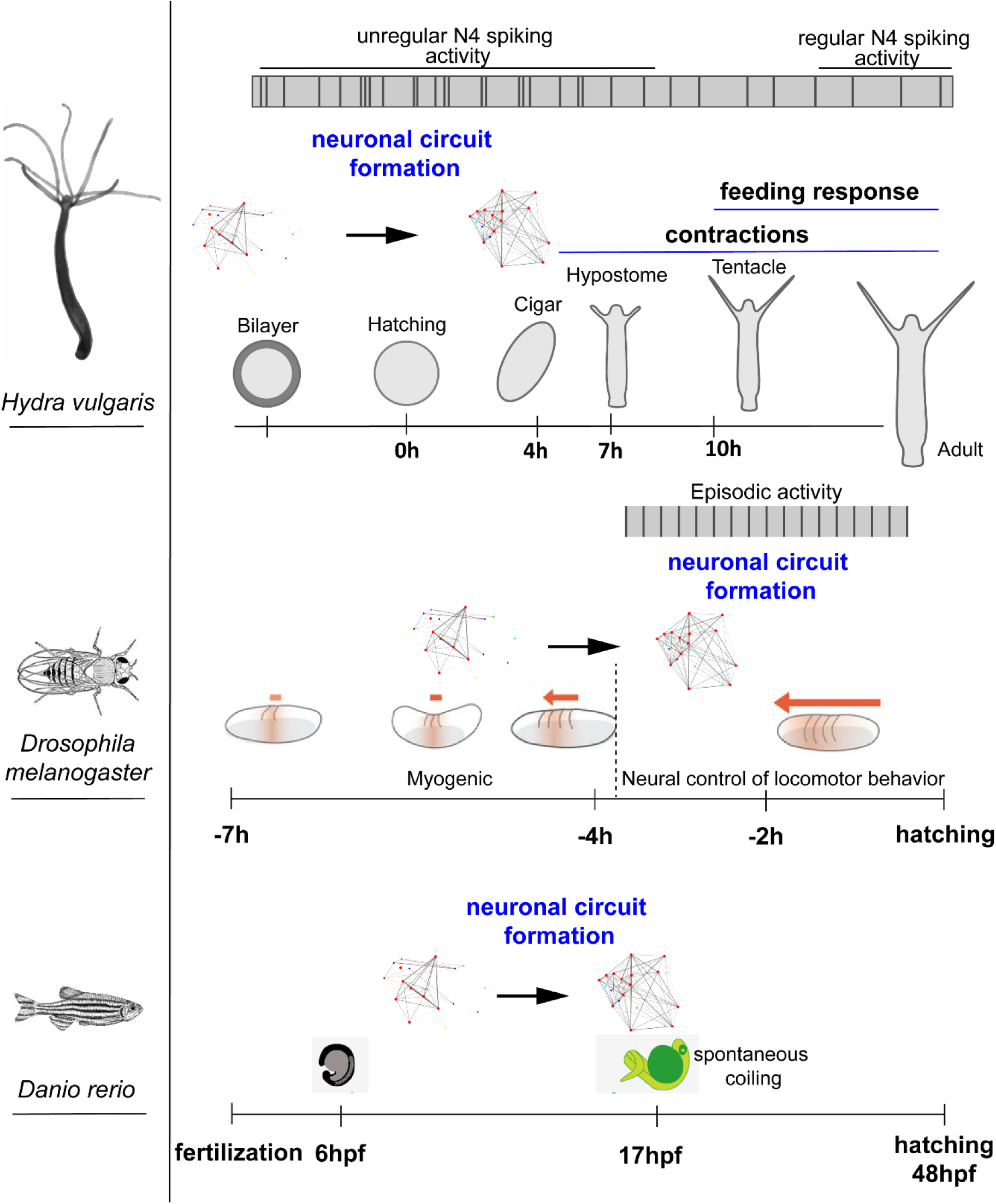
Neuronal circuit formation is universal and emerges in the absence of a centralized information processing.

## Methods

### Animals

#### Cultivation

Hydra polyps (*Hydra vulgaris* AEP) were cultivated in standard *Hydra* medium (CaCl_2_ 0.042g/l; MgSO_4_ x 7H_2_O 0.081g/l; NaHCO_3_ 0.042g/l; K_2_CO_3_ 0.011g/g in dH_2_O) according to previous established protocols^52^. The animals were cultured in either 20 x 20 x 6cm culture dishes or 250ml glass beakers at 18°C depending on culture size. Polyps were held at a 12/12h light cycle and strictly fed three times a week to induce sexual reproduction, utilizing *Artemia nauplii.* Animals were washed after a digestion period of 8-14h. Adult polyps were not fed 48h prior to any experiments. For long-term calcium recordings polyps were fed at day 2 and day 8 after removing the cuticle.

### Embryos

#### Collecting

*Hydra* cultures were frequently observed and embryos were collected in batches of 25 and stored in 6-well plates (Eppendorf cat#0030720113) at 18°C and 12/12h light conditions in *Hydra* medium and observed daily (from day 10 on) with a dissecting microscope (Wild M.3) throughout their development. To determine the hatching rate, embryos were cut off from the female polyp 24 hpf during the cuticle stage and collected in 6-well plates.

#### Determination time points

To distinguish between important changes in morphology, up to three late non-hatched embryos (day 10+) were recorded using a dissecting stereomicroscope (Olymups SZX16) equipped with a camera (Olympus DP74) at 1 frame per minute (Olympus cellSense Standard 1.18) in a microscope slide with a depth of 5mm and covered with a cover glass to avoid evaporation of the *Hydra* media.

#### Preparation

Embryos and hatchlings were collected at specific developmental stages, relaxed with ice-cold 2% Urethane in *Hydra* media and fixed in Zamboni for 2h (RT) or overnight (4°C). Bilayer-staged embryos were opened with a scalpel or a preparation needle to remove the cuticle prior to fixation. Next, the samples were washed three times for 15min in PBS with 0.1% Tween (PBST) before storing them for further analysis in PBST + NaN_3_ at 4°C to avoid mold formation.

#### Generating germ-free embryos

To generate germ-free embryos, cuticle-staged embryos (24 hpf) were used, which were cut off from female *Hydra vulgaris* AEP cultures. Embryos were treated with an antibiotic cocktail containing Ampicillin (50mg/ml), Neomycin (50mg/ml), Rifampicin (50mg/ml in DMSO), Streptomycin (50mg/ml) and Spectinomycin (60mg/ml) as described previously^53^. The embryos were kept under dark conditions in sterile cups for 3 days without changing the antibiotic solution. After 3 days embryos were transferred to freshly prepared antibiotic solution under sterile conditions and incubated another 3 days. Throughout the 6-day treatment, embryos were incubated at 18°C on an orbital shaker (IKA, Rocker 3D digital cat#0004001000) at 12 rpm After a total of 6 days, embryos were rinsed with sterile HM to avoid transfer of antibiotic residue and placed in sterile media afterwards. To ensure the absence of bacteria, sterility tests were performed at 2- and 8-days post-antibiotic treatment. Thus, two embryos of each batch were homogenized in 100μl sterile media and plated on R2A agar plates and incubated at 18°C. To further ensure the germ-free status we tested via PCR by isolating whole DNA from single embryos (Qiagen cat#69504) using universal 16S rRNA Primers^54^. Absent amplification products and colonies after 1 week of incubation on the R2A agar confirmed the germ-free status.

#### Recolonization experiments

Germ-free embryos were generated as described above to perform recolonization experiments. Two days post-antibiotics treatment, the sterile embryos were transferred to a glass beaker (Carl Roth cat#C095.1) along with 70 wild-type adult polyps to restore the. Cultures with adult polyps and embryos were fed two times a week and washed 12-14h after feeding. Embryos were observed two times a day after 10d post-cuticle formation.

### Transgenic lines

The transgenic line N4 (GCaMP6s) used to generate embryos has been published in another study^12^. Here, animals in the F1 generation were used to obtain transgenic embryos, thus ensuring an incorporation of the construct. To generate a pan-neuronal cross-breeding line for Ca^2+^ activity (GCaMP6s) we only used animals from F1-generations of previously published lines^6,12^ (N4, N6, alpha tubulin). For cross-breeding, alpha tubulin females were incubated with N4 males. Positive embryos were cultured and sexual reproduction was induced. Next, these animals were used to cross-breed females with N6 males to generate the P46 line.

### Histology

#### Immunohistochemistry

Previously fixed embryos were washed 3 x 15min with PBST, permeabilized for 30min in PBS + 0.5% Triton X-100 followed by a blocking step (PBST + 1% bovine serum albumin, BSA) for 1h. Animals were then incubated overnight at 4°C with a primary antibody in PBST + 1% BSA. Primary antibodies used in this study were: anti-FMFR (BMA Biomedicals, cat# T-4322, 1:1000), anti-GLWamide (KPNAYKGKLPIGLW-amide, 1:2000, Takahashi 2003) and anti-GFP (Biozol, cat# GFP-1010, 1:1000). After incubation with the primary antibody, four 15min washing steps (PBST + 1% BSA) were conducted before adding the secondary antibody. For this study, the following secondary antibodies were used: goat anti-chicken IgY Alexa Fluor 488 (Invitrogen, cat# A11039, 1:1000) and goat anti-rabbit IgG (Invitrogen, cat# A11034). Samples were incubated for 2h (RT) without light in the antibody solution followed by another four washing steps in PBST + 1%BSA (with 0.5% Tween). Samples were either stored at 4°C for a short period or mounted in Mowiol on a coverslip (Carl Roth, cat# 1871.2) covered by a cover glass (Carl Roth, cat# 1870.2) separated by an imaging spacer (Grace Bio-Labs, cat#SS1X20) and stored in a humidity chamber at 4°C until imaging.

#### Histology sections

Samples were embedded in epoxy resin as described for TEM ultrastructural analysis. Semi-thin (1 µm) epoxy resin sections were cut on a Leica UC7 ultramicrotome using a histo diamond knive. Semi-thin sections were deposited on Superfrost^®^ Plus adhesive slides, stained with Richardson’s solution (alkaline solution of azure II (50%) and methylene blue (50%)), embedded between a glass slide and a cover slip using a CV Mount medium (Leica) and imaged using a Zeiss LSM900 microscope equipped with a 20x Plan-Apochromat objective (NA 0.8), an AxioCam 305 color camera and a ZEN 3.2 system including automated tiling function.

#### Ultrastructural analysis by transmission electron microscopy

Hydra embryos and hatchlings were relaxed with 2% urethane prepared in Hydra medium for 1 min and transferred to 10mM cacodylate buffer + 1% glutaraldehyde for fixation. Polyps were post-fixed with 1% osmium tetroxide prepared in 1.5% potassium ferricyanide/dH_2_O for 1 h on ice, and contrasted en-bloc with 2% aqueous uranyl acetate for 1 h at room temperature. Dehydration was performed at room temperature with ascending ethanol series (each step 15 min), followed by acetone (2 × 30 min). After progressive infiltration with epoxy resin (Epon_812 substitute, Sigma), samples were heat polymerized. Thin, 80-nm sections were prepared using Leica UC7 ultramicrotome and a Diatome diamond knife, and contrasted with saturated uranyl acetate and Reynolds lead citrate. Sections were imaged with a Tecnai G2 Spirit BioTWIN transmission electron microscope (FEI), operated at 80 kV, and equipped with a LaB6 filament, a CCD side-mounted MegaView III G2 camera, and iTEM v.5 software (both Olympus Soft Imaging solutions) as described previously^6^.

### Imaging and analysis

The fixed and stained embryos and hatchlings were imaged with a confocal laser-scanning microscope (LSM900, Zeiss) using the Airyscan mode an a Plan-Apochromat 20x / NA 0.8 objective, or the Axio Vert.A1 (Zeiss) using Colibri 7 (Zeiss) as a light source. Images were further processed with Zen Blue 3.4 software (Zeiss) or Fiji^55^. For determination of total neuron numbers, all RFamide-pos neurons were counted only in the head area of the embryos and hatchlings with a minimum of five samples for each stage and treatment. In addition, to identify neuronal connections we analyzed neurites originated from a neuron soma for up to 10 cells per sample with a minimum amount of 5 samples for each stage. To examine neuronal numbers and neurites, composites of stacks (maximum projections) were used, whereas brightness and contrast were modified to increase visibility. The analysis was carried out using R (2023.12.0+369)^56^ over RStudio IDE. To visualize we used the plugins “ggplot” (v3.5.1), “tidyr” (v1.3.1), “ggpubr” (v0.6.0) and “dplyr” (v1.1.4).

### In vivo single-cell Ca^2+^ imaging

To analyze the neuronal activity on single-cell resolution, we developed a self-made chamber which allows for examining the whole embryo throughout long-term recordings at following temperatures: standard: 18°C; cold: 13°C; warm: 23°C. To adjust the temperature, an air-conditioned room was used, which was set to the requested temperature 3 days prior to recordings. Embryos in the Bilayer stage were placed into an imaging spacer affixed to a 24x50mm cover slip (Carl Roth cat#1871.2) filled with *Hydra* media and closed with a small 22x22mm cover slip (Carl Roth, cat#H874.2) with two openings on both sides of the imaging spacer to fill and refill media. To reopen the chamber, the upper plastic liner remained on the spacer which constituted a standard height of 0.24mm during recordings. Hatchlings and adult polyps were placed into commercially available channel slides that were 0.1 or 0.2mm in height (ibidi cat#80661, ibidi cat#80161). Once the samples were placed into the chambers or channel slides, animals were allowed to adapt for 10min plus 2min to the laser prior to recordings. For our experiments a 14% laser intensity was used and animals during the first 7.5h were recorded for 30min with 25fps. After recording, the samples rested for 60min and remained within the self-made chamber, which was filled with an additional amount of 400µl *Hydra* media to slightly lift the cover glass to increase the space between top and bottom. After a resting time of 60min, the previously added media was removed and recorded as previously described. Animals after 1 or 10 days were recorded for 10min with the same recording setup. Imaging was performed using the Axio Vert.A1 (Zeiss) using Colibri 7 (Zeiss) as light source equipped with a 38HE filter (Zeiss), the Axiocam 705 mono (Zeiss) and a 10x Plan Apo objective. The recordings were implemented to Zen Blue 3.4 software (Zeiss) for image export, further processed in ImageJ (Source) to 8-bit and in total 2000 frames were selected from each dataset which were analyzed with ICY software^57^. Tracked neurons were automatically traced using the protocol “Detection (with cluster un-mixing) and Tracking of neurons with emc2” whereas parameters were constituent adjusted depending on individual samples. Accuracy was manually perused for each tracked neuron, missing links connected and merged. Non-specific and false tracked were removed.

### Community Detection

Community detection results were obtained using the Leiden algorithm^27^ that was shown to outperform the previously state-of-the-art Louvain algorithm. The resolution parameter for the constant Potts model (CPM) used as the quality measure of the Leiden algorithm was set to *γ* = 0.96. Connectivity data to be processed by the algorithm were given by correlation matrices estimated from the measurement and simulation data. Since the CPM mode of the Leiden algorithm failed to process the complete correlation matrices, only non-negative values were used. To increase the accuracy of the correlation analysis, the measurement were processed with the spike inference method MLspike^58^. This allowed us to remove the drifting baseline and the noise present in the measured fluorescence trace and therefore normalized recorded tracks to improve the comparison.

### Electrical circuit model

#### Neuron model

The circuit model builds upon a modified Morris-Lecar circuit as a neuron model^59^, because it is mathematically less complex than the Hodgin-Huxley model, but still biologically accurate. The exploited modified model replaced the calcium channel of the original Morris-Lecar model by a sodium channel, and added an additional calcium channel for enabling the calculation of a corresponding calcium concentration. The modified Morris-Lecar circuit is described by

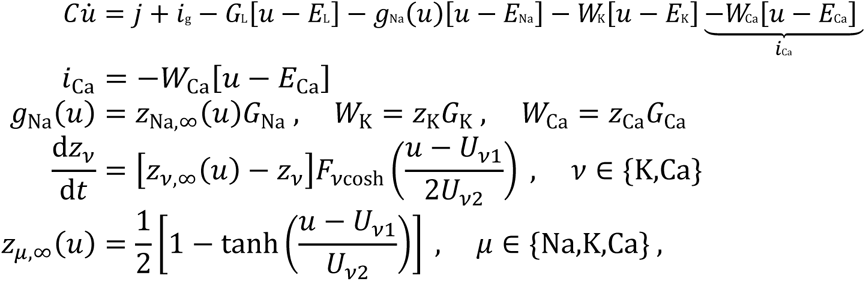

where *u* is the membrane potential, *j* is an external current, *i*_Ca_ is the calcium current, and *i*_g_ is the current received from other neurons. *g*_Na_(*u*), *W*_K_, and *W*_Ca_ are functions for the channel opening of the specific ion channels. *z*_nu_ describes the fraction of open ion channels, while *z*_mu*,∞*_ models the opening and closing process. Parameter values are listed in Tab. 1.

**Table 1:** Parameters for the circuit model.

| Morris-Lecar parameters |  |  |
| --- | --- | --- |
| Input current amplitude | $j =$ | 220 nA |
| Capacitance | $C =$ | 8 μF |
| Sodium channel conductance | $G_{Na} =$ | 20 mS |
| Sodium resting potential | $E_{Na} =$ | 50 mV |
| Sodium channel threshold voltage | $U_{Na1} =$ | 5 mV |
| Sodium channel edge steepness | $U_{Na2} =$ | 13 mV |
| Potassium channel conductance | $G_K =$ | 30 mS |
| Potassium resting potential | $E_K =$ | -90 mV |
| Potassium channel threshold voltage | $U_{K1} =$ | 0 mV |
| Potassium channel edge steepness | $U_{K2} =$ | 10 mV |
| Potassium channel opening rate | $F_K =$ | 7.84 Hz |
| Calcium channel conductance | $G_{Ca} =$ | 4.6 nS |
| Calcium resting potential | $E_{Ca} =$ | 111 mV |
| Calcium channel threshold voltage | $U_{Ca1} =$ | -1 mV |
| Calcium channel edge steepness | $U_{Ca2} =$ | 12 mV |
| Calcium channel opening rate | $F_{Ca} =$ | 500 Hz |
| Gap junction parameters |  |  |
| Leakage channel conductance | $G_L =$ | 20 μS |
| Leakage resting potential | $E_L =$ | -70 mV |
| Minimum gap junction conductance | $Wg,0 =$ | 14.7 μS |
| Maximum gap junction conductance | $Wg,1 =$ | 15 μS |
| Gap junction slope of change | $S_g =$ | 0.38 Hz |
| Leakage memristor retention characteristic | $Sg,ret =$ | 190 mHz |
| Positive threshold voltage for gap junction memristor | $Ug,p =$ | 50 mV |
| Negative threshold voltage for gap junction memristor | $Ug,n =$ | -150 mV |
| RC circuit parameters |  |  |
| Capacitance | $C_\eta =$ | 0.1 mF |
| Resistance | $R_\eta =$ | 7.9 kΩ |
| Voltage for resting calcium concentration | $U_{\eta,rest} =$ | 48.16 nV |

#### Calcium concentration model

We generated an artificial calcium concentration that is physically linked to the experimentally observed fluorescence traces, by integrating the calcium current of the modified Morris-Lecar circuits^60^. Based on previous studies^58^, we achieved this by using an additional RC circuit governed by

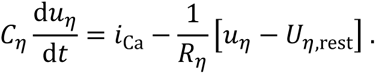

Here, *u_η_* represents the calcium concentration *η*, and *U_η,_*_rest_ represents the resting calcium con-centration *η*_rest_. Parameter values are given in Tab. 1.

#### Model for neuronal interconnections

We assumed that neurons in the population N4 are coupled via gap junctions. This is based on the observation that neurons of N4 in adult polyps show strong synchronous behavior. For the circuit model, we represented gap junctions between the neurons by nonlinear resistors with memory, in short memristors. These memristors allowed us to incorporate the growth of neuronal connections into the circuit model. We can assume that the resistance value of the space between the contact points of neurons depends on the distance. In this sense, two distant neurons exhibit a high resistance value, while a connection via gap junctions exhibits a low resistance value. The growing connection can therefore, in a very simplistic way, be modeled by gradually decreasing the resistance value. We implemented this via the equations

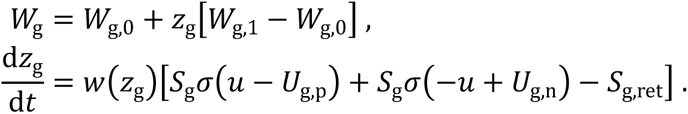

Here, *z*_g_ is an internal state variable reflecting how distant the contact points of two neurons are, such that *z*_g_ = 0 implies very distant neurons and *z*_g_ = 1 indicates a connection via a gap junction. *w*(·) is a window function that ensures *z*_g_ ∈ [0, 1], and *σ*(·) is the Heaviside function. Parameters are given in Tab. 1.

A dynamic network growth with variable number of neurons is achieved as follows: New neurons are added to the network by setting the external currents *j* for the not yet considered neurons to 0A, such that only the considered ones are active. Moreover, memristors *W*_g_ are added only between the active neurons.

### Vector-valued circuit model

We represented the complete circuit model for the 30 coupled neuron models in a vector-valued fashion, since this allowed for a more compact representation that simplified the simulation approach. The vector-valued model is governed by

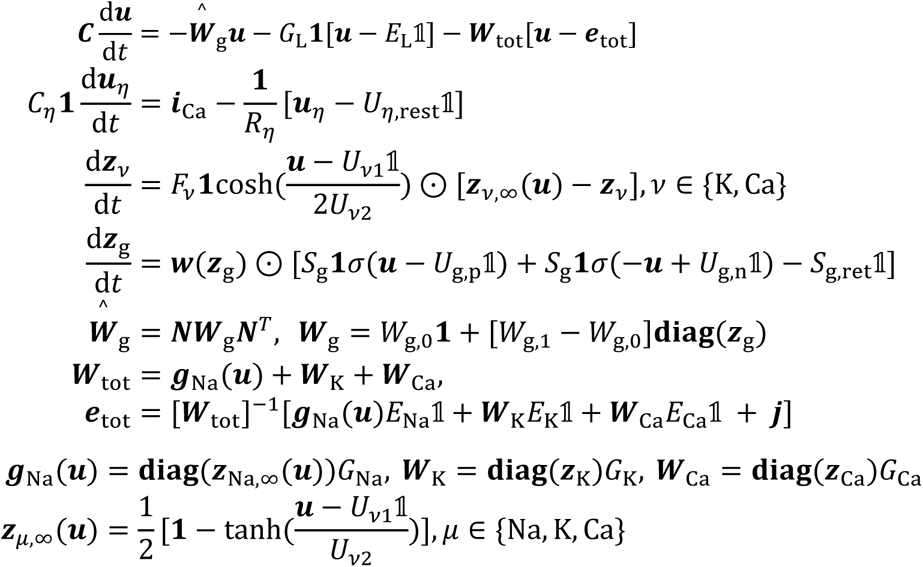

with ***j****, **u**, **i***_Ca_*, **u**_η_, **z***_Ca_*, **z***_Ca_*, **z***_K_ ∈ R^30*×*1^, and ***z***_g_ ∈ R^435*×*1^. The interconnection structure of the neuron circuits is described by the incidence matrix **N** ∈ R^30*×*435^, **1** is the unity matrix, 𝟙 is a vector of ones, ⊙ is the Hadamard product, and **diag**(·) converts its vector argument into a diagonal matrix for the capacitances. Note that parameters of the neuron model such as ***C*** are a diagonal matrix ∈ R^30*×*30^. We assumed identical parameters for every neuron, since specific electrophysiological characterizations of *Hydra* neurons are not available.

### Wave digital simulation

Electrical circuits were translated into wave digital models^61^ by first decomposing them into their one- and multiports and their interconnection structure. Ports and their electrical quantities were then discretized, for which the trapezoidal rule was used as numerical integration method that is especially required for reactive elements such as capacitors. Resulting ports and intercon-nection structures were finally translated into their wave digital counterparts using the bijective transformation

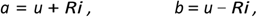

with the incoming wave *a*, the reflected wave *b*, and the port resistance *R >* 0. This yields a wave digital model than can be programmed and therefore be exploited for emulating the original circuit model. We applied this technique to the vector-valued circuit model that resulted in the corresponding wave digital model shown in Fig.XXX of the supplementary material. Note that this included artificial delay elements called iterator elements used to solve implicit relationships^59,62,63^

### Statistics and data analysis

All statistics were performed using R and R-Studio as IDE as well as MatLab and Python. We tested for different assumptions for the different tests. Further, we tested outcomes of values as non-parametric tests (Kruskal-Wallis, Dunn test) and p-values were adjusted with the Bonferroni method. The replicate number (n) for each dataset is indicated in the figures and/or the figure legends as well as the statistical method. Unless specified differently, values are presented as median ± standard deviation. Statistical significance was depicted as asterisks when p<0.05. (* p<0.05; ** p<0.01; *** p<0.001)

## Supporting information

SupplVid1_Hatching-stage

SupplVid2_Cigar-stage

SupplVid3_Tentacle-stage

SupplVid4_Adult-polyp

SupplVid5_artificial_calcium-imaging

## Data availability

Data used in this study are presented in the Supplementary Table or available and shared by the corresponding author upon request

## Code availability

The used codes and scripts to reproduce the main experiments of this study implemented in Python, Matlab and R are available at GitHub (https://github.com/ChNoack-Ki/Hydra-Embryos)

## Acknowledgments

We thank Claus C. Hilgetag and Alexander Klimovich for discussions and critical reading of the manuscript. We greatly appreciate excellent technical support and microinjection of the *Hydra* embryos by Jörg Wittlieb. We further thank Katja Duwe-Schrinner for the preparation and finalization of the graphical abstract. We are grateful for excellent assistance of Ulrike Voigt (Central Microscopy, Kiel University) in preparing resin sections for TEM analysis. The Anti-GLWamide antibody was kindly provided by Toshitaka Fujisawa. This work was funded by the German Research Foundation (Deutsche Forschungsgemeinschaft), the CRC 1182 “Origin and Function of Metaorganisms” (to TCGB and M.B., project C1 and Z2), the CRC 1461 “Neurotronics: Bio-inspired Information Pathways” (Project-ID 434434223 – SFB 1461; TPA04; to TCGB, KHO and AK) and DFG Grant KL3475/2-1 (to AK). TCGB gratefully appreciates support from the Canadian Institute for Advanced Research. We acknowledge funding for the Zeiss LSM 900 with Airyscan microscope by the German Research Foundation: INST 257/650-1 FUGG.

## Contributions

C.N., C.G. and T.C.G.B conceptualized the project. C.N. designed experiments. C.N. performed neural activity experiments. C.N., S.J. and C.G. analyzed activity experiments. S.J. developed observer model. C.N., O.M. and L.-M.H performed and analyzed Immunohistochemistry and microbiota experiments. O.M. implemented germ-free assay for embryos. J.W. generated transgenic line. U.R. and M.B. performed electron microscopy and subsequent data analysis. C.N., S.J. and T.C.G.B. wrote the manuscript.

## Ethics declarations

The authors declare no competing interests.

## References

1. Richter, L. M. & Gjorgjieva, J. Understanding neural circuit development through theory and models. Current Opinion in Neurobiology vol. 46 39–47 Preprint at 10.1016/j.conb.2017.07.004 (2017).

2. Arendt, D. Elementary nervous systems. Philosophical Transactions of the Royal Society B: Biological Sciences 376, (2021).

3. Giez, C., Klimovich, A. & Bosch, T. C. G. Neurons interact with the microbiome: An evolutionary-informed perspective. Neuroforum vol. 27 89–98 Preprint at 10.1515/nf-2021-0003 (2021).

4. Klimovich, A. et al. Prototypical pacemaker neurons interact with the resident microbiota. doi:10.1073/pnas.1920469117/-/DCSupplemental.

5. Cazet, J. F. et al. A chromosome-scale epigenetic map of the Hydra genome reveals conserved regulators of cell state. Genome Res 33, 283–298 (2023).

6. Giez, C. et al. Satiety controls behavior in Hydra through an interplay of pre-enteric and central nervous system-like neuron populations. Cell Rep 43, (2024).

7. Keramidioti, A. et al. A new look at the architecture and dynamics of the Hydra nerve net. Elife 12, (2024).

8. Dupre, C. & Yuste, R. Non-overlapping Neural Networks in Hydra vulgaris. Current Biology 27, 1085–1097 (2017).

9. Siebert, S. et al. Stem cell differentiation trajectories in Hydra resolved at single-cell resolution. Science (1979) 365, (2019).

10. Rg Wittlieb, J., Khalturin, K., Lohmann, J. U., Anton-Erxleben, F. & Bosch, T. C. G. Transgenic Hydra Allow in Vivo Tracking of Individual Stem Cells during Morphogenesis. www.pnas.orgcgidoi10.1073pnas.0510163103 (2006).

11. Klimovich, A., Wittlieb, J. & Bosch, T. C. G. Transgenesis in Hydra to characterize gene function and visualize cell behavior. Nat Protoc 14, 2069–2090 (2019).

12. Giez, C. et al. Multiple neuronal populations control the eating behavior in Hydra and are responsive to microbial signals. Current Biology 33, 5288–5303.e6 (2023).

13. Lovas, J. R. & Yuste, R. Ensemble synchronization in the reassembly of Hydra’s nervous system. Current Biology 31, 3784–3796.e3 (2021).

14. Tomczyk, S. et al. Loss of Neurogenesis in Aging Hydra. Dev Neurobiol 79, 479–496 (2019).

15. Yamamoto, W. & Yuste, R. Peptide-driven control of somersaulting in Hydra vulgaris. Current Biology 33, 1893–1905.e4 (2023).

16. Kim, S., Badhiwala, K. N., Duret, G. & Robinson, J. T. Phototaxis is a satiety-dependent behavioral sequence in Hydra vulgaris. Journal of Experimental Biology (2024) doi:10.1242/jeb.247503.

17. Luo, L. Architectures of neuronal circuits. Science vol. 373 Preprint at 10.1126/science.abg7285 (2021).

18. Dale Purves, George J. Augustine & David Fitzpatrick. Neuroscience. (Sinauer Associates, Sunderland, MA, USA, 2011).

19. Donald O. Hebb. *The Organization of Behavior*. (John Wiley & Sons, New York, NY, USA, 1949).

20. Irwin B. Levitan & Leonard K. Kaczmarek. The Neuron. (Oxford University Press Inc., New York, NY, USE, 2015).

21. György Buzsáki. The Brain from Inside Out. (Oxford University Press, New York, NY, USA, 2019).

22. George L. Gerstein, Marc J. Bloom & Pedro E. Maldonaldo. Central Auditory Processing And Neural Modeling. (Springer, New York, NY, USA, 1998).

23. Löwel, S. & Singer, W. Löwel-Singer_1992_Selection of Intrinsic Horizontal Connections in the.

24. Eytan, D. & Marom, S. Dynamics and effective topology underlying synchronization in networks of cortical neurons. Journal of Neuroscience 26, 8465–8476 (2006).

25. Khalturin, K. et al. Transgenic stem cells in Hydra reveal an early evolutionary origin for key elements controlling self-renewal and differentiation. Dev Biol 309, 32–44 (2007).

26. Klimovich, A. V. & Bosch, T. C. G. Rethinking the Role of the Nervous System: Lessons From the Hydra Holobiont. BioEssays vol. 40 Preprint at 10.1002/bies.201800060 (2018).

27. Traag, V. A., Waltman, L. & van Eck, N. J. From Louvain to Leiden: guaranteeing well-connected communities. Sci Rep 9, (2019).

28. Augustin, R. et al. A secreted antibacterial neuropeptide shapes the microbiome of Hydra. Nat Commun 8, 1–8 (2017).

29. He, J. & Bosch, T. C. G. Hydra’s Lasting Partnership with Microbes: The Key for Escaping Senescence? Microorganisms 10, (2022).

30. Domin, H. et al. Sequential host-bacteria and bacteria-bacteria interactions determine the microbiome establishment of Nematostella vectensis. Microbiome 11, (2023).

31. Murillo-Rincon, A. P. et al. Spontaneous body contractions are modulated by the microbiome of Hydra. Sci Rep 7, (2017).

32. Santiago Ramon y Cajal. The Croonian Lecture. - La Fine Structure Des Centres Nerveux. (The Royal Society, London, UK, 1894).

33. Bang, S. et al. Engineered neural circuits for modeling brain physiology and neuropathology. Acta Biomaterialia vol. 132 379–400 Preprint at 10.1016/j.actbio.2021.06.024 (2021).

34. Martínez, D. E. & Bridge, D. Hydra, the everlasting embryo, confronts aging. International Journal of Developmental Biology 56, 479–487 (2012).

35. Peter Robin Hiesinger. Self-Assembling Brain. (Princeton University Press, 2021).

36. Ochs, K. & Jenderny, S. An equivalent electrical circuit for the Hindmarsh-Rose model. International Journal of Circuit Theory and Applications 49, 3526–3539 (2021).

37. Jenderny, S., Ochs, K. & Xue, D. A memristive circuit for self-organized network topology formation based on guided axon growth. Sci Rep 14, (2024).

38. Taubenheim, J. et al. Bacteria-and temperature-regulated peptides modulate β-catenin signaling in Hydra. doi:10.1073/pnas.2010945117/-/DCSupplemental.

39. Bourn, J. J. & Dorrity, M. W. Degrees of freedom: temperature’s influence on developmental rate. Current Opinion in Genetics and Development vol. 85 Preprint at 10.1016/j.gde.2024.102155 (2024).

40. Fraune, S. & Bosch, T. C. G. Long-Term Maintenance of Species-Specific Bacterial Microbiota in the Basal Metazoan Hydra. www.pnas.org/cgi/content/full/ (2007).

41. Fraune, S. et al. In an early branching metazoan, bacterial colonization of the embryo is controlled by maternal antimicrobial peptides. Proc Natl Acad Sci U S A 107, 18067–18072 (2010).

42. Franzenburg, S. et al. Bacterial colonization of Hydra hatchlings follows a robust temporal pattern. ISME Journal 7, 781–790 (2013).

43. Bathia, J., Miklos, M., Gyulai, I., Fraune, S. & Tokolyi, J. Environmental microbial reservoir influences the Hydra-associated bacterial communities. Preprint at 10.21203/rs.3.rs-4881820/v1 (2024).

44. Vuong, H. E. et al. The maternal microbiome modulates fetal neurodevelopment in mice. Nature 586, 281–286 (2020).

45. Coulson, B. et al. Critical periods in Drosophila neural network development: Importance to network tuning and therapeutic potential. Frontiers in Physiology vol. 13 Preprint at 10.3389/fphys.2022.1073307 (2022).

46. Marín, O. Developmental timing and critical windows for the treatment of psychiatric disorders. Nature Medicine vol. 22 1229–1238 Preprint at 10.1038/nm.4225 (2016).

47. Lee, E., Lee, J. & Kim, E. Excitation/Inhibition Imbalance in Animal Models of Autism Spectrum Disorders. Biological Psychiatry vol. 81 838–847 Preprint at 10.1016/j.biopsych.2016.05.011 (2017).

48. Sadeqzadeh, E., De Bock, C. E. & Thorne, R. F. Sleeping Giants: Emerging Roles for the Fat Cadherins in Health and Disease. Med Res Rev 34, 190–221 (2014).

49. Kohsaka, H., Okusawa, S., Itakura, Y., Fushiki, A. & Nose, A. Development of larval motor circuits in Drosophila. Development Growth and Differentiation vol. 54 408–419 Preprint at 10.1111/j.1440-169X.2012.01347.x (2012).

50. Nelson, J. C. & Granato, M. Zebrafish behavior as a gateway to nervous system assembly and plasticity. Development (Cambridge) 149, (2022).

51. Humberto R. Maturana & Francisco J. Varela. *The Tree of Knowledge*. (Shambhala, Boston, MA, USA, 1992).

52. Bode, H., et al. Quantitative Analysis of Cell Types during Growth and Morphogenesis in Hydra. Wilhelm goux’ Archly vol. 171 (1973).

53. Franzenburg, S. et al. Distinct antimicrobial peptide expression determines host species-specific bacterial associations. Proc Natl Acad Sci U S A 110, (2013).

54. Weisburg, W. G., Barns, S. M., Pelletier, D. A. & Lane, D. J. 6S Ribosomal DNA Amplification for Phylogenetic Study. JOURNAL OF BACTERIOLOGY vol. 173 https://journals.asm.org/journal/jb (1991).

55. Schindelin, J., et al. Fiji: An open-source platform for biological-image analysis. Nature Methods vol. 9 676–682 Preprint at 10.1038/nmeth.2019 (2012).

56. RC Team. R Core Team R: A Language and Environment for Statistica Computing. Preprint at (2020).

57. Lagache, T., Hanson, A., Pérez-Ortega, J. E., Fairhall, A. & Yuste, R. EMC ^2^ : A versatile algorithm for robust tracking of calcium dynamics from individual neurons in behaving animals. Preprint at 10.1101/2020.06.22.165696 (2020).

58. Deneux, T. et al. Accurate spike estimation from noisy calcium signals for ultrafast three-dimensional imaging of large neuronal populations in vivo. Nat Commun 7, (2016).

59. Jenderny, S. & Ochs, K. Wave digital model of calcium-imaging-based neuronal activity of mice. International Journal of Numerical Modelling: Electronic Networks, Devices and Fields 36, (2023).

60. Maravall, M., Mainen, Z. F., Sabatini, B. L. & Svoboda, K. Estimating intracellular calcium concentrations and buffering without wavelength ratioing. Biophys J 78, 2655–2667 (2000).

61. Fettweis, A. Wave Digital Filters: Theory and Practice.

62. Schwerdtfeger, T. & Kummert, A. Nonlinear Circuit Simulation by Means of Alfred Fettweis’ Wave Digital Principles.

63. Jenderny, S., Ochs, K. & Hövel, P. Power consumption during forward locomotion of C. elegans: an electrical circuit simulation. European Physical Journal B 97, (2024).

