## Supplementary Figures for "Assembly of a functional neuronal circuit in embryos of an ancestral metazoan is influenced by environmental signals including the microbiome"

**Extended Data Fig.1: Stage-specific maturation of *Hydra* hatchlings and neurons.**

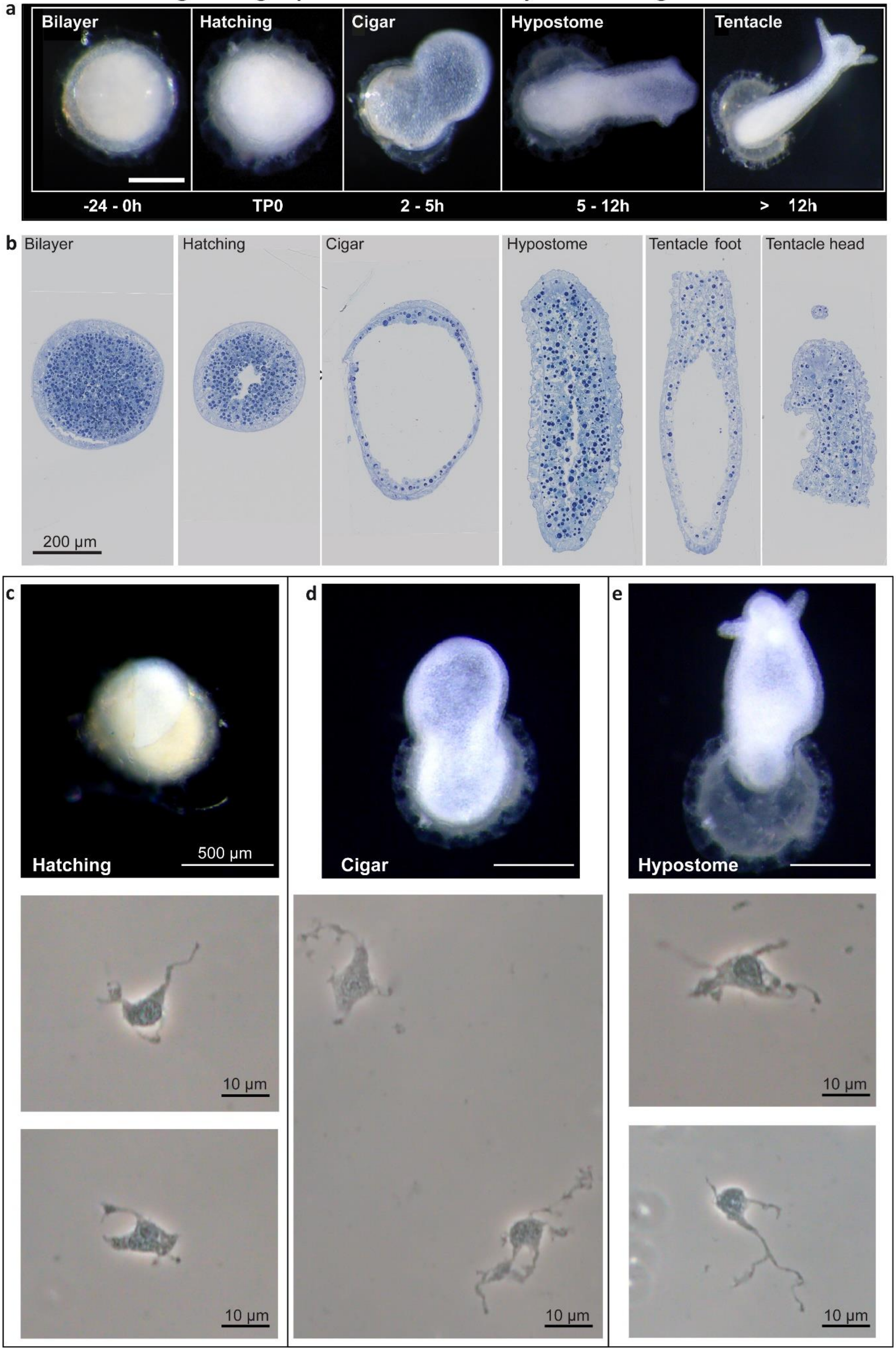

**a,** Determination timepoints based on morphology changes during development of *Hydra* embryos and hatchlings. First accessible samples during Bilayer stage, lower caption indicate time needed until or after rupture of the cuticle. Scale bars, 500 µm. **b,** Longitudinal resin sections of each stages colored with Richardson stain. **c,** Currently hatching *Hydra* polyp (top). Macerations of single individuals during Hatching stage revealed neuronal cells with short neurites, low number of branches and with a minor amount of heterochromatin (middle on bottom pictures) compared to further developed individuals. **d,e,** Later-staged hatchlings in contrast display extended neurites and more branches as well as a higher amount of heterochromatin.

### Extended Data Fig.2: Potential communication via EDVs and gap junctions.

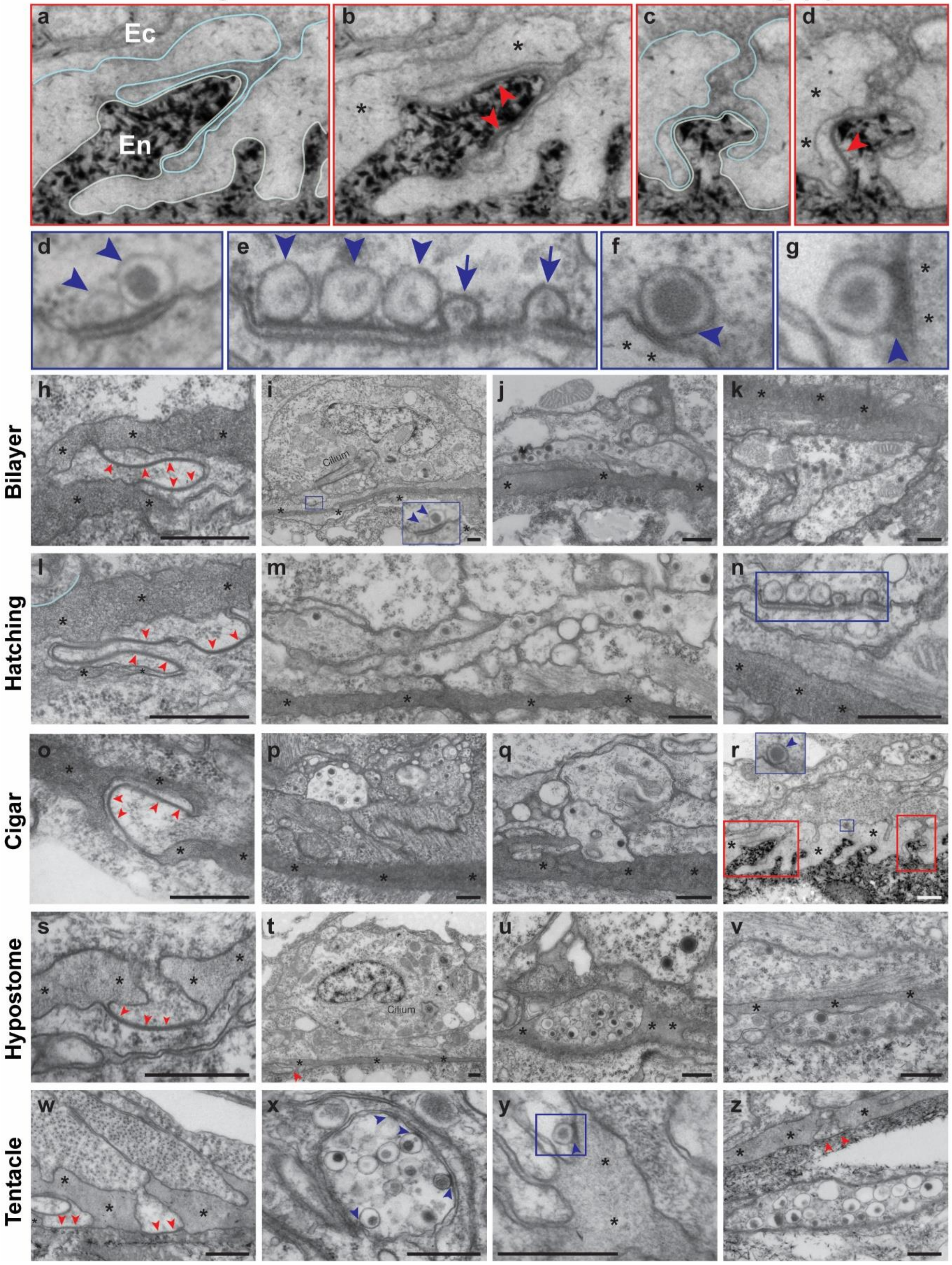


**a-d,** Magnifications from cellular extensions with gap-junctions (red arrowheads) of ectodermal (Ec) and endodermal (En) cells interact across the mesoglea (asterisks) in the hypostome region (Cigar stage) referring to **r.** The cytoplasm of the endodermal cells is contaminated with uranyl acetate. Light blue line: ectodermal cell plasma membrane; light green line: endodermal plasma membrane. **d,e,** Electron-dense core vesicles (EDV) in neurons docked to the plasma membrane referring to **i** and **n**. **f,g,** EDV in epithelial cells of ectoderm, docked to the plasma membrane that is exposed to the mesoglea. **h-z,** Pictures presented in the main Fig.2 without coloring. Scale bars 500nm.

### Extended Data Fig.3: Cell-cell connections.

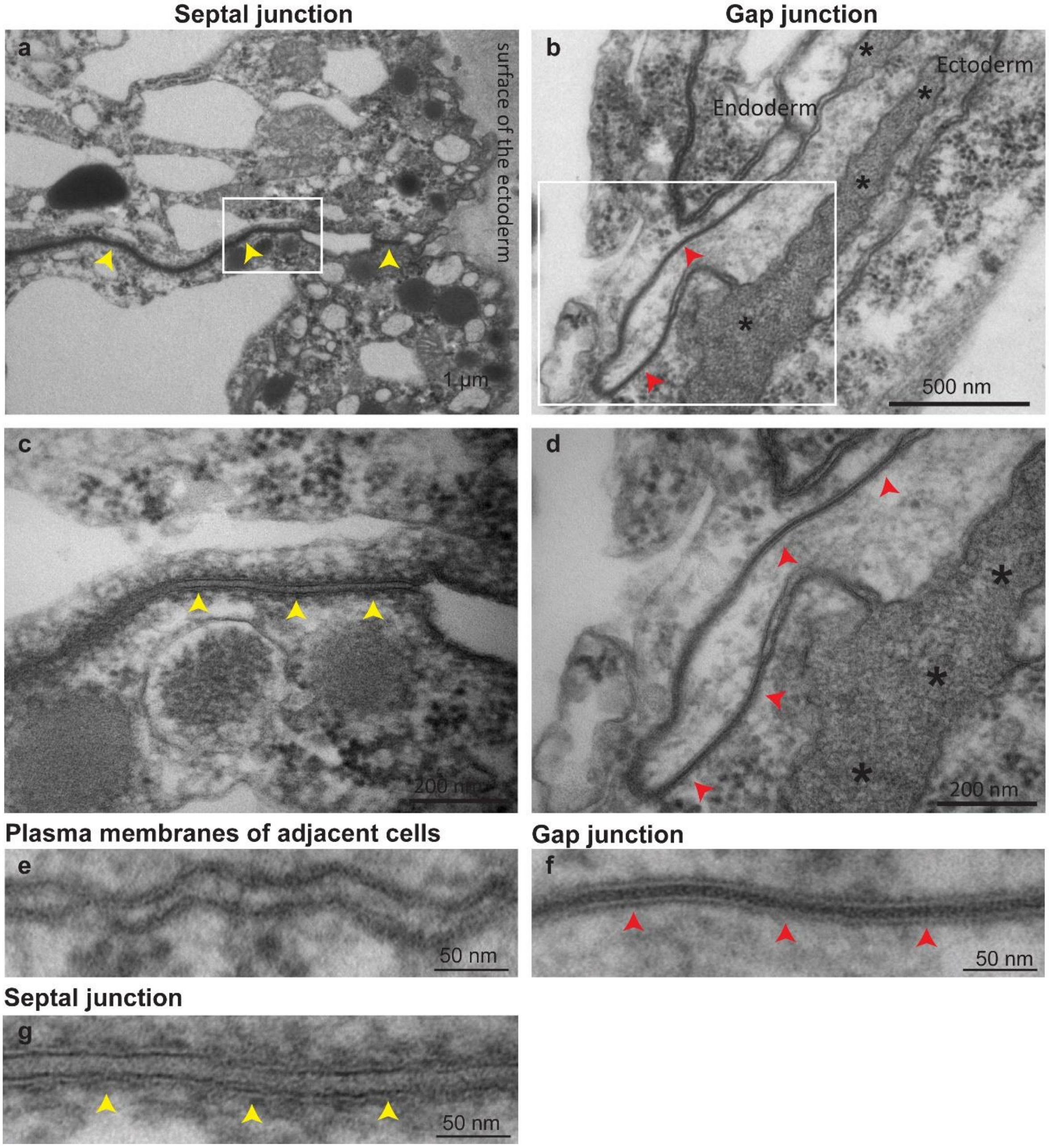


**a-g,** Enlarged structures focusing on plasma membranes and different junctions found in early *Hydra* hatchlings. Red arrowheads: gap-junctions; yellow arrowheads: septal-junctions.

### Extended Data Fig.4: Schematic overview of findings with TEM analysis.

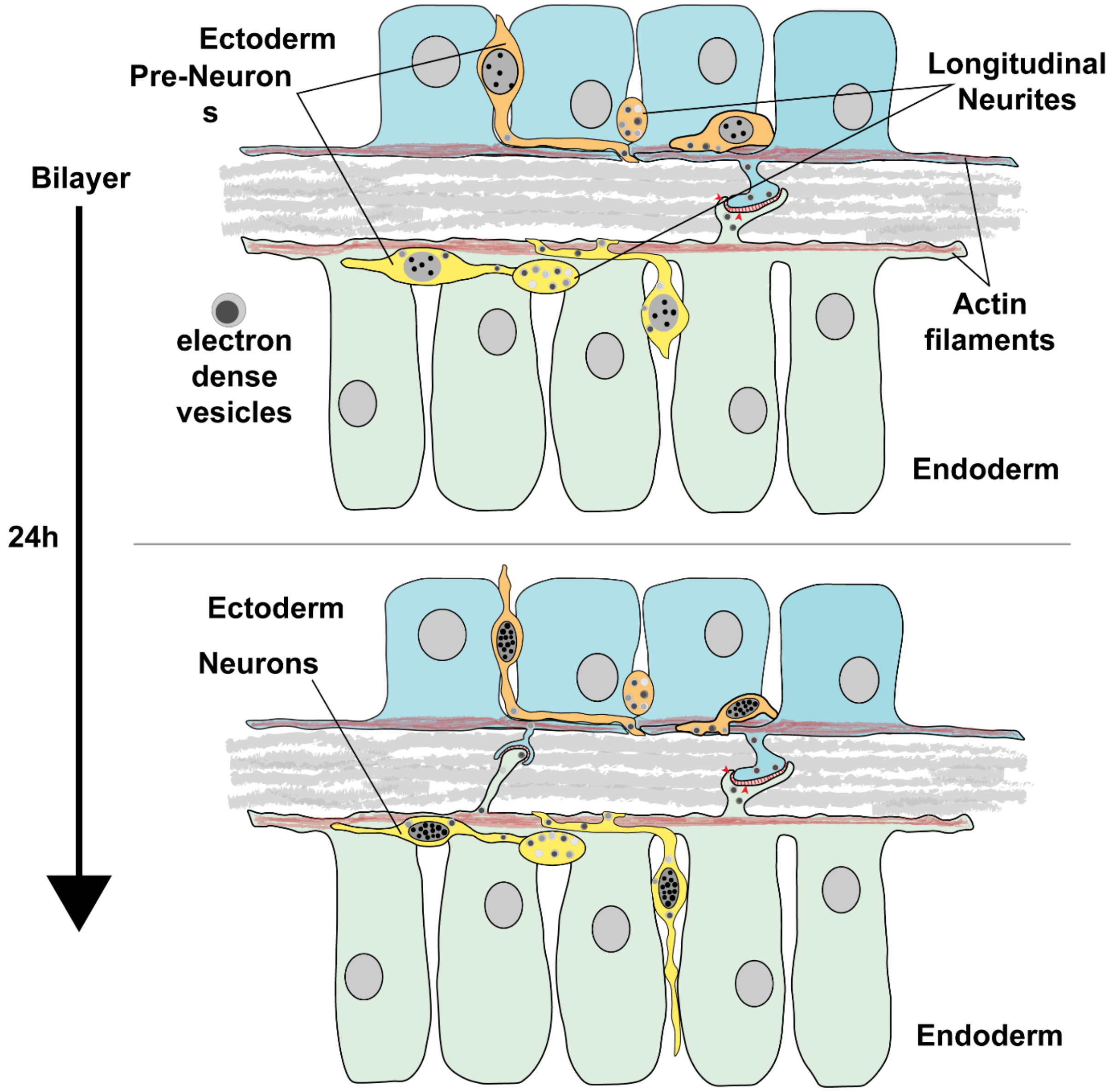


Schematic representation of the core findings in the TEM analysis starting in the Bilayer stage and ending in the Tentacle stage, covering a developmental timeframe of approximately 24h. Over the course of time we observed a maturation of neurons as well as an increase of connections between endodermal and ectodermal epithelial cells throughout the mesoglea. Gap-junctions as well as electron-dense vesicles appear to drive and support a rapid development of the nervous system. TEM analysis revealed a widespread communication potential between neurons and mesoglea, neurons and epithelial cells, epithelial cells and mesoglea as well as among endodermal and ectodermal epithelial cells.

**Extended Data Fig.5: Calcium activty and antibody stainings revealed a concealed pool. of inactive or non-mature neurons.**

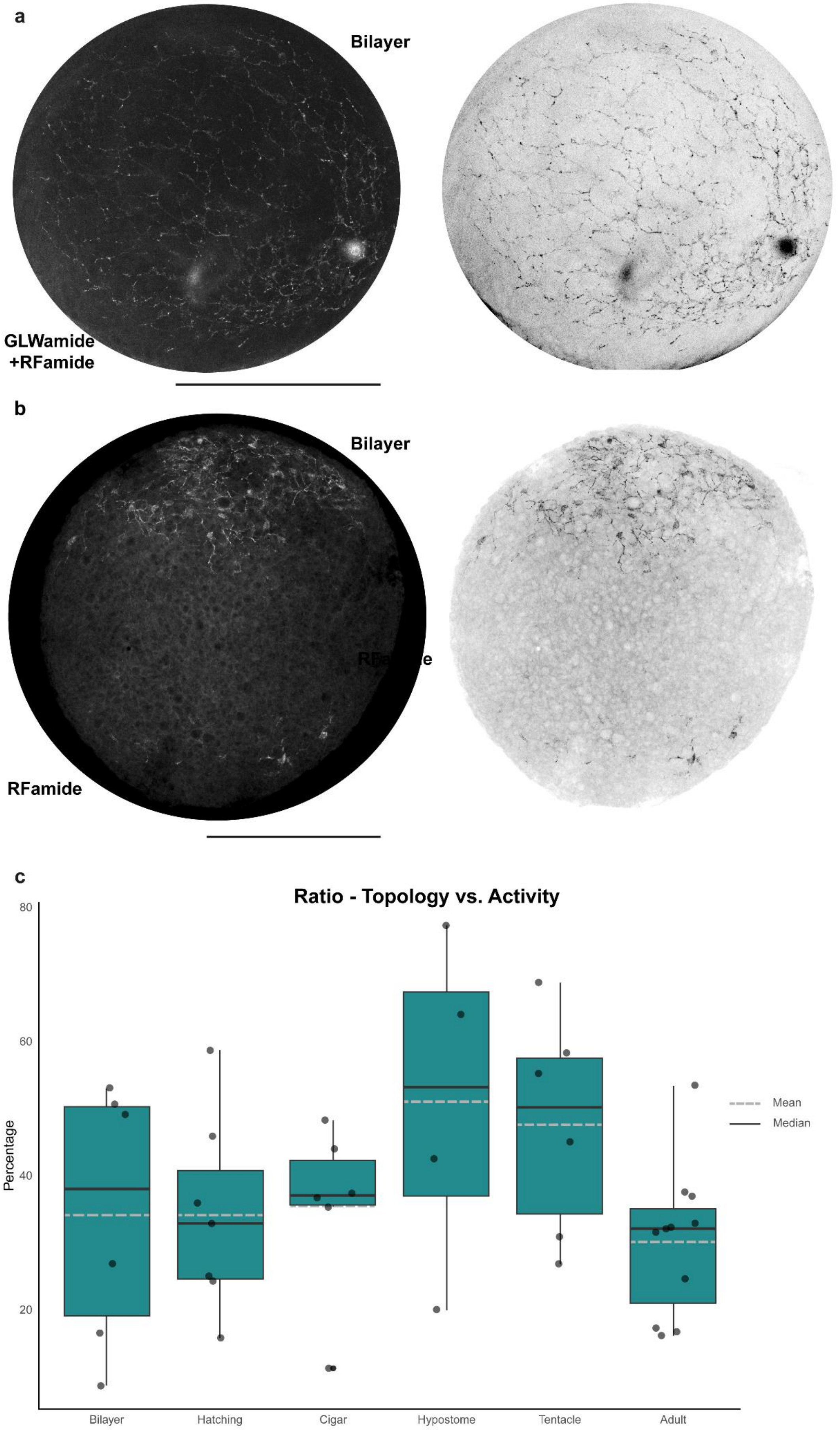

**a,** Immunohistochemistry against GLWamide and RFamide revealed already established connections between neuronal structures during Bilayer stage (left: monochrome; right: inverted). **b,** Immunohistochemistry against RFamide exhibited an early polarization even in bilayer-staged embryos, as specific RFamide-stained populations are exclusively expressed in the future head or foot area. **c,** Comparison of the ratio between topology and activity indicated a declining number of stained and $Ca^{2+}$ active neurons. Considering the Hypostome stage about 50% of cells belong equally to either active or stained neurons suggested an integration of cells needed for the nerve net architecture facilitating a rapid assembly (n = 4-11). Adult polyps displayed a ratio of 30% of active neurons.

### Extended Data Fig.6: Ca2+ activity N4-hatchlings.

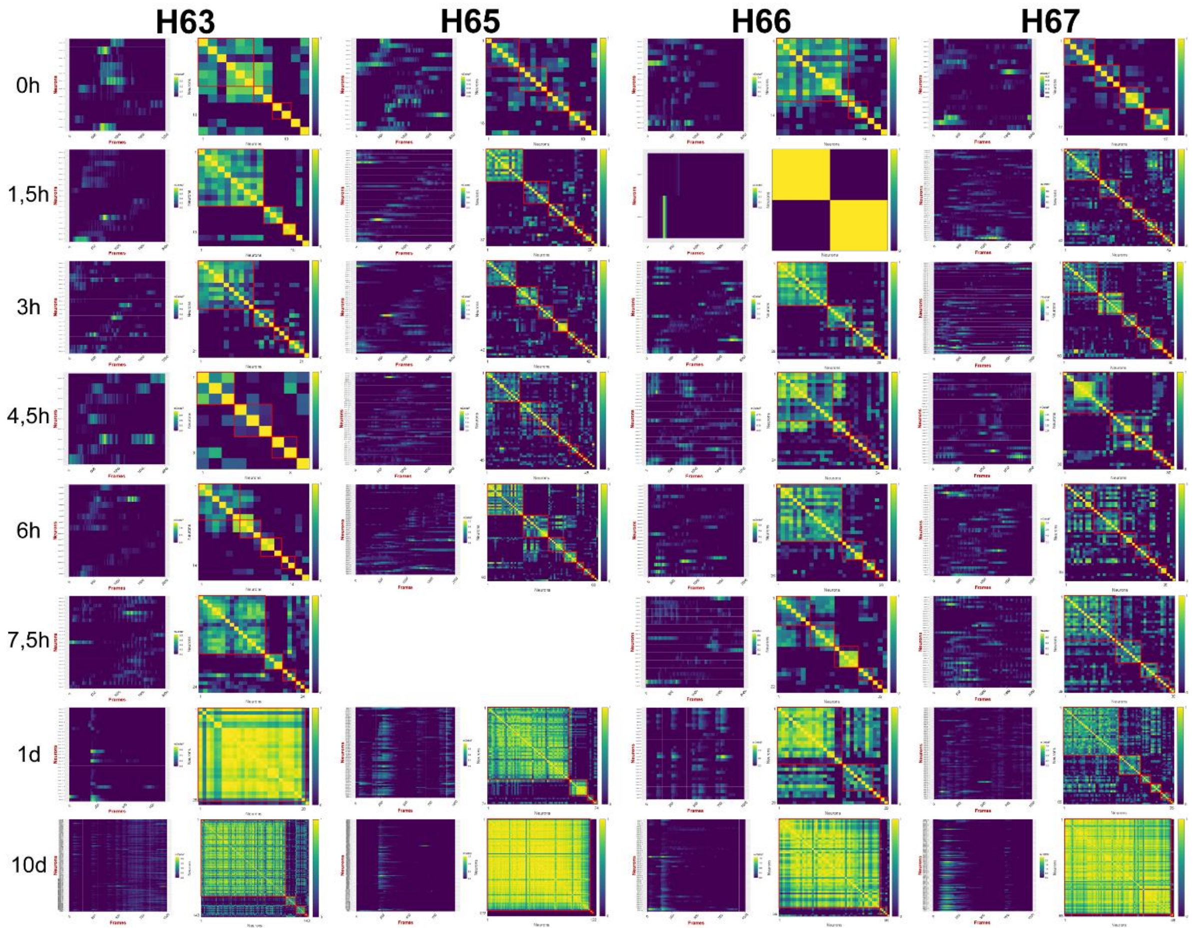


Entire overview of single-cell tracking in *Hydra* hatchlings at defined timepoints. Synchronization level and community detection in hatchlings recorded under standard conditions (18°C) referring to **Fig. 4a,b**. (n=5)

**Extended Data Fig.7: Entire structure of the assembling network.**

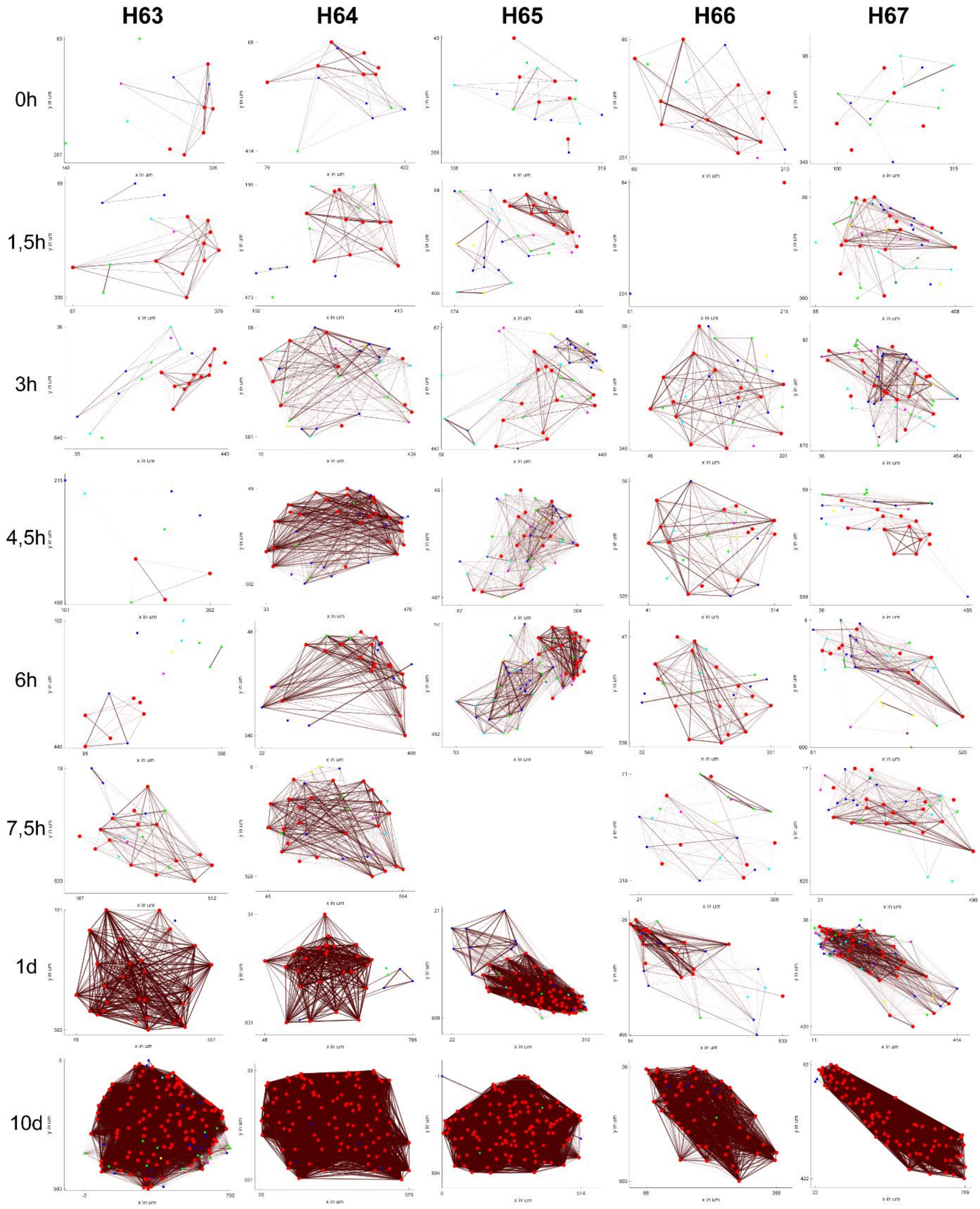


Overview of nerve net assembly at defined timepoints under standard conditions (18°C) based on position data. (n = 5)

**Extended Data Fig.8: Circuit model and additional simulations.**

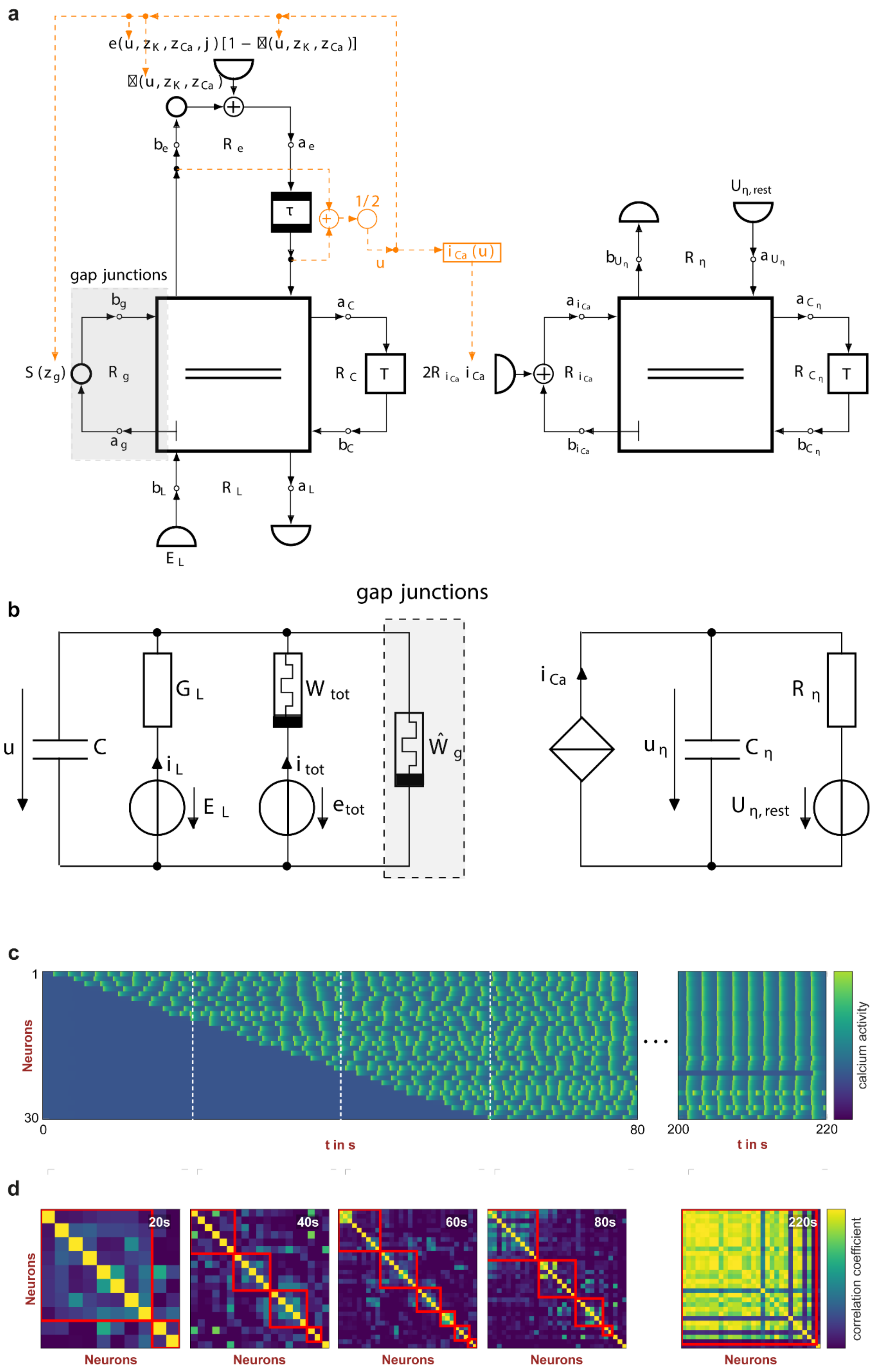

**a,** Electrical circuit diagram for the vector-valued circuit model described in the methods section. **b,** Wave digital flow diagram corresponding to the vector-valued circuit model. **c,d,** Emulation results for higher frequency variations exhibited by the deployed neuron model based on observation in *Hydra* hatchlings recorded under warm (23°C) conditions. **c,** Simulated calcium activity. **d,** Detected communities for a simulated nerve net with identical neuron parameters are highlighted by red squares.

**Extended Data Fig.9: Network assembly cold (13°C) temperatures.**

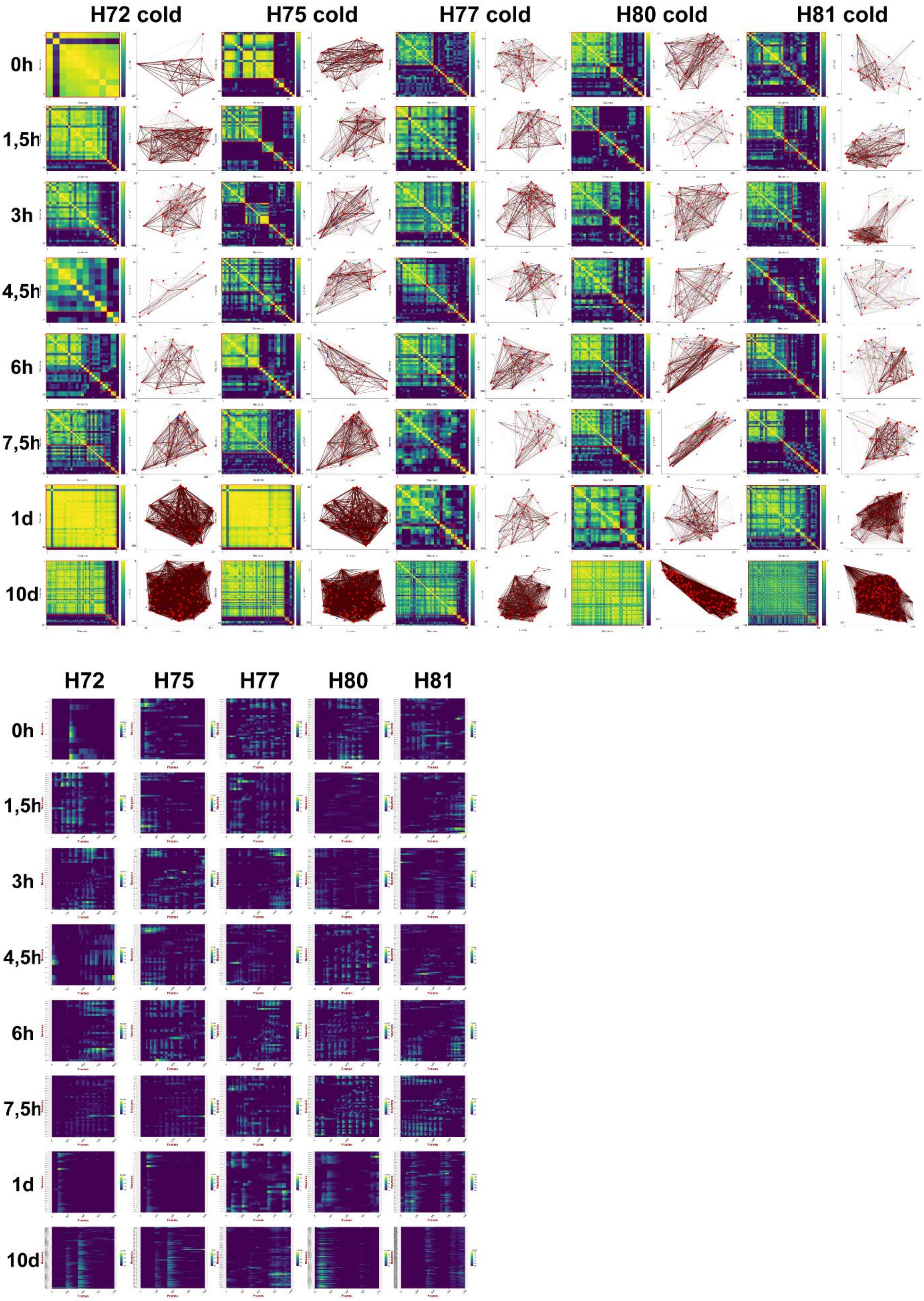


Entire overview of $Ca^{2+}$ recordings under cold (13°C) conditions. Illustrated here, community detection, network assembly based on position data and synchronization plots throughout all recording timepoints. (n = 5)

**Extended Data Fig.10: Network assembly warm (23°C) temperatures.**

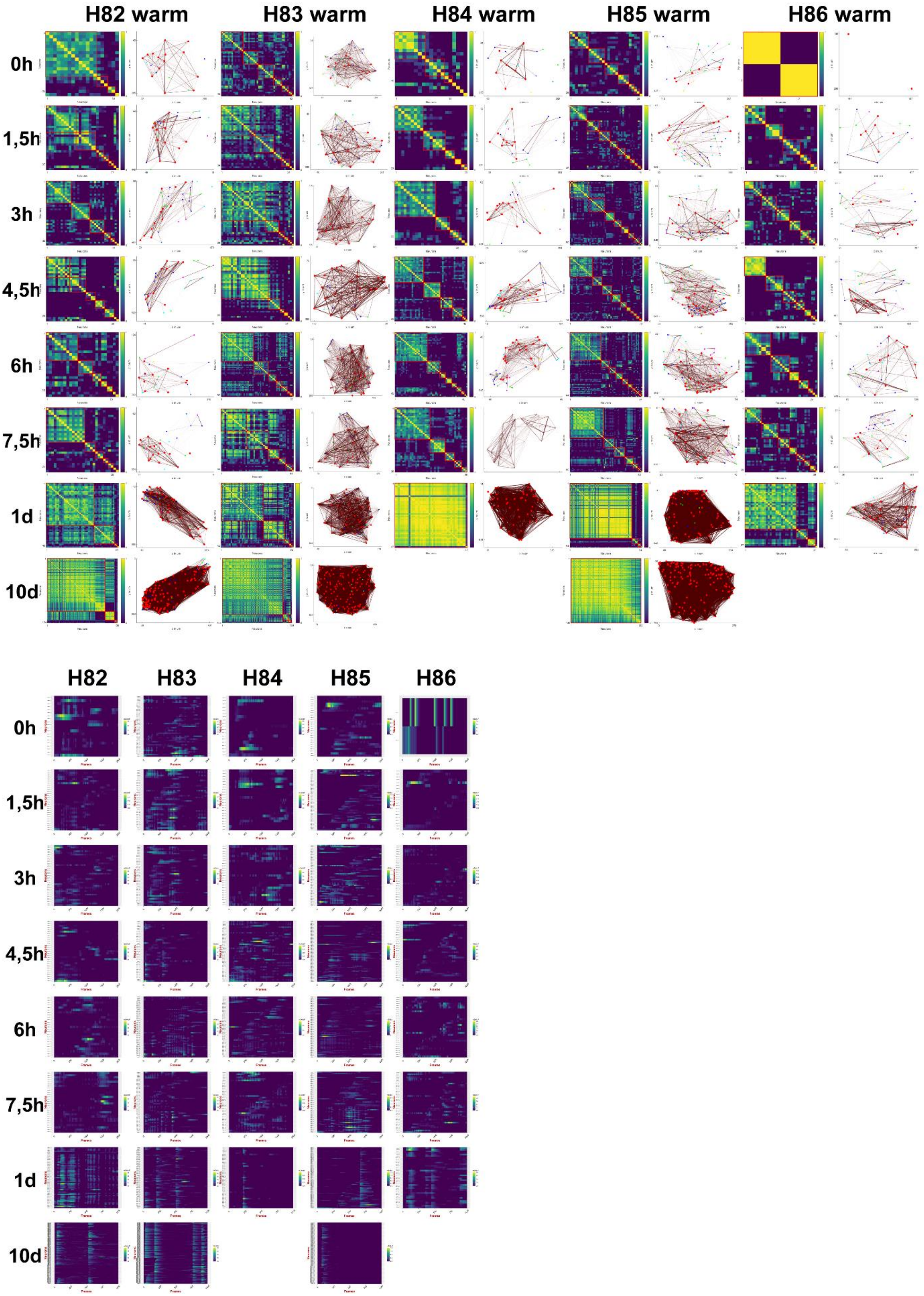


Entire overview of $Ca^{2+}$ recordings under warm (23°C) conditions. Illustrated here, community detection, network assembly based on position data and synchronization plots throughout all recording timepoints. (n = 5)

### Extended Data Fig.11: Immunohistochemistry of microbial impact.

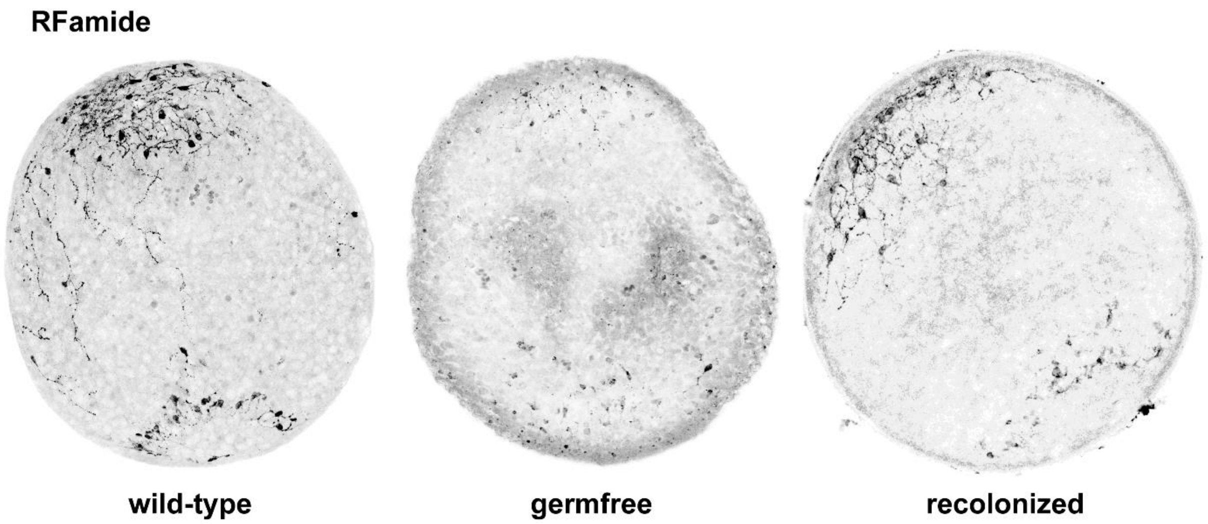


Immunohistochemistry of hatchlings during Hatching stage against RFamide, corresponding to **Fig.7 j-l**.
